# Rab35 Governs Apicobasal Polarity Through Regulation of Actin Dynamics During Sprouting Angiogenesis

**DOI:** 10.1101/2022.01.13.476231

**Authors:** Caitlin R. Francis, Hayle Kincross, Erich J. Kushner

**Affiliations:** Department of Biological Sciences, University of Denver, Denver, CO

**Keywords:** angiogenesis, lumenogenesis, blood vessel development, actin, Rab35, DENNd1c, trafficking, apical membrane, cytoskeleton, zebrafish

## Abstract

In early blood vessel development, trafficking programs, such as those using Rab GTPases, are tasked with delivering vesicular cargo with high spatiotemporal accuracy. However, the function of many Rab trafficking proteins remain ill-defined in endothelial tissue; therefore, their relevance to blood vessel development is unknown. Rab35 has been shown to play an enigmatic role in cellular behaviors which differs greatly between tissue-type and organism. Importantly, Rab35 has never been characterized for its potential contribution in sprouting angiogenesis; thus, our goal was to map Rab35’s primary function in angiogenesis. Our results demonstrate that Rab35 is critical for sprout formation; in its absence apicobasal polarity is entirely lost in vitro and in vivo. To determine mechanism, we systematically explored established Rab35 effectors and show that none are operative in endothelial cells. However, we find that Rab35 partners with DENNd1c, an evolutionarily divergent guanine exchange factor, to localize to actin. Here, Rab35 regulates actin polymerization, which is required to setup proper apicobasal polarity during sprout formation. Our findings establish that Rab35 is a potent regulator of actin architecture during blood vessel development.

## INTRODUCTION

Angiogenesis is the process of sprouting and growth of new blood vessels from preexisting ones and is the primary driver of network expansion [1–4]. Many extrinsic and intrinsic biological systems have been shown to affect endothelial biology and, by extension, blood vessel formation. Membrane trafficking is one such system that is less well-characterized in endothelial tissue, but has recently become more appreciated as additional organotypic trafficking signatures are aligned with important endothelial behaviors [5–8]. Membrane trafficking refers to vesicular transport of protein(s) to, or in vicinity of, the plasma membrane [9–11]. Here, trafficking regulators, such as Rab GTPases, interface with a host of effectors involved in receptor recycling, cytoskeletal regulation, shunting to degradative organelles, lumen formation, basement membrane secretion, and many other signaling events [9, 12, 13]. Indeed, critical to the understanding of how endothelial cells build dynamic and resilient vascular structures is the regulation of membrane trafficking during angiogenic development.

The GTPase Rab35 has been shown to be a multi-faceted regulator of membrane trafficking and continues to be an intensely researched Rab family member [14]. The promiscuity of Rab35 touching multiple pathways has created a cognitive bottleneck in attempting to assign function in any system, due to its seemingly endless diversity of roles. For instance, Rab35 has been shown to be involved in cytokinesis as well as transcytosis of the apical protein podocalyxin during lumen biogenesis in epithelial cysts [15, 16]. In other investigations, Rab35 has been reported to be a negative regulator of the integrin recycling protein Arf6 via its effector ACAP2 [17–19]. Additionally, MICAL1 has been shown to also facilitate Rab35’s association with Arf6 and play a role in actin turnover [19–21]. In drosophila, Rab35 regulates apical constriction during germband extension as well as actin bundling via recruitment of fascin [22, 23]. To date, there is no unified study on Rab35 taking into account its many disparate functions in any tissue. Regarding blood vessel function, no endothelial studies exist detailing how, or if, Rab35 functions in sprouting angiogenesis.

In the current study, our goal was to comprehensively characterize Rab35’s role in sprouting angiogenesis. To do so, we took a holistic approach in investigating established partners of Rab35 and characterized their effect on sprouting behaviors and downstream cellular morphodynamics in vitro and in vivo. Primarily using a 3-dimensional sprouting assay, our results revealed that Rab35 is required for sprouting as its loss significantly disrupts apicobasal polarity. Focusing on Rab35 effectors, we demonstrate that of the many reported effectors only ACAP2 was capable of directly binding Rab35 in endothelial cells. However, upon investigating ACAP2 and its target Arf6, we determined this established Rab35 trafficking cascade was largely insignificant with regard to sprouting angiogenesis. Excluding all other pathways, we focused on the Rab35 guanine exchange factor (GEF), DENNd1c, and its role in localizing Rab35 to actin structures. Our results demonstrate that DENNd1c facilitates Rab35 tethering to the actin cytoskeleton. Once on actin, Rab35 acts as a positive regulator of actin polymerization and is critical for formation of proper actin architecture. In vivo, we show the requirement of Rab35 in zebrafish blood vessel development using a gene editing approach. Overall, our results provide novel evidence of a focused role for Rab35 as a regulator of actin assembly during sprouting angiogenesis.

## MATERIALS AND METHODS

### Reagents

All reagent information is listed in the reagents table in the supplementary information.

### Cell Culture

Pooled Human umbilical vein endothelial cells (HUVECs) were purchased from PromoCell and cultured in proprietary media (PromoCell Growth Medium, ready-to-use) for 2-5 passages. All cells were maintained in a humidified incubator at 37°C and 5% CO_2_. Small interfering RNA (ThermoFisher) was introduced into primary HUVEC using the Neon® transfection system (ThermoFisher). Scramble, Rab35, Podocalyxin, ACAP2, OCRL, MICAL-L1, DENNd1a, DENNd1b, and DENNd1c were purchased from (ThermoFisher) and resuspended to a 20µM stock concentration and used at 0.5 µM. Normal human lung fibroblasts (NHLFs, Lonza) and HEK-A (ThermoFisher) were maintained in Dulbeccos Modified Medium (DMEM) supplemented with 10% fetal bovine serum and antibiotics. Both NHLFs and HEKs were used up to 15 passages. For 2-dimensional live-imaging experiments, cells were imaged for one minute at baseline before treatment with CK-666 (1μM), and then imaged for an additional two minutes using 5 second intervals. For ligand-modulated antibody fragments tether to the mitochondria (Mito-LAMA) experiments procedures were carried out as previously described [47]. Briefly, cells were electroporated with mito-LAMA (pCDNA3.0_mitoLAMA-G97), the protein of interest, as well as the target of interest (tag-RFP-DENND1c, mCherry-Arp2, or LifeAct-647). Baseline images were taken for 1 minute (5 second intervals), treated with Trimethoprim (TMP, 500uM) and promptly imaged for an additional 5 minutes at 5 second intervals.

### Sprouting Angiogenesis Assay

Fibrin-bead assay was performed as reported by Nakatsu et al. 2007 [25]. Briefly, HUVECs were coated onto microcarrier beads (Amersham) and plated overnight. SiRNA-treatment or viral transduction was performed the same day the beads were coated. The following day, the EC-covered microbeads were embedded in a fibrin matrix. Once the clot was formed media was overlaid along with 100,000 NHLFs. Media was changed daily along with monitoring of sprout development. Sprout characteristics were quantified in the following manner. Sprout numbers were determined by counting the number of multicellular sprouts (sprouts that did not contain at least 3 cells were not used in the analysis) emanating from an individual microcarrier beads across multiple beads in a given experiment. Sprout lengths were determined by measuring the length of a multicellular sprout beginning from the tip of the sprout to the microcarrier bead surface across multiple beads. Percent of non-lumenized sprouts were determined by quantifying the proportion of multicellular sprouts whose length (microcarrier bead surface to sprout tip) was less than 80% lumenized across multiple beads. Sprout widths were determined by measuring the sprout width at the midpoint between the tip and the microcarrier bead across multiple beads. Actin accumulation were defined by actin puncta with a diameter greater than 1.5μm. Experimental repeats are defined as an independent experiment in which multiple cultures, containing numerous sprouting beads were quantified; this process of quantifying multiple parameters across many beads and several cultures was replicated on different days for each experimental repeat.

### Plasmid Constructs

The following constructs were procured for this study: GFP-Rab35 S22N inactive (gift from Peter McPherson; Addgene plasmid # 47426); GFP-Rab35 WT (gift from Peter McPherson, Addgene plasmid # 47424); GFP_Rab35 Q67L (gift from Peter McPherson, Addgene plasmid # 47425); mEmerald-Fascin-C-10 (gift from Michael Davidson, Addgene plasmid # 54094); pARF6(Q67L)-CFP (gift from Joel Swanson, Addgene plasmid # 11387); pARF6(T27N)-CFP (gift from Joel Swanson, Addgene plasmid # 11386); pARF6-CFP (gift from Joel Swanson, Addgene plasmid # 11382); pcDNA3-HA-human OCRL (gift from Pietro De Camilli, Addgene plasmid # 22207); pCDNA3.0_mitoLAMA-G97 (gift from Kai Johnsson, Addgene plasmid # 130705); pGST1-GGA3-VHS (gift from James Hurley, Addgene plasmid # 44420); mEmerald-ARP2-C-14 (gift from Michael Davidson, Addgene plasmid # 53992); MICAL-L1 (Origene, RG214051); and DENNd1c (Origene, RC206410);

### Lentivirus and Adenovirus Generation and Transduction

Lentivirus was generated by using the LR Gateway Cloning method [24]. Genes of interest and fluorescent proteins were isolated and incorporated into a pME backbone via Gibson reaction [69]. Following confirmation of the plasmid by sequencing the pME entry plasmid was mixed with the destination vector and LR Clonase. The destination vector used in this study was pLenti CMV Neo DEST (705-1) (gift from Eric Campeau & Paul Kaufman; Addgene plasmid #17392). Once validated, the destination plasmids were transfected with the three required viral protein plasmids: pMDLg/pRRE (gift from Didier Trono; Addgene plasmid # 12251), pVSVG (gift from Bob Weinberg; Addgene plasmid #8454) and psPAX2 (gift from Didier Trono; Addgene plasmid #12260) into HEK 293 cells. The transfected HEKs had media changed 4 hours post transfection. Transfected cells incubated for 3 days and virus was harvested.

Adenoviral constructs and viral particles were created using the Adeasy viral cloning protocol (9). Briefly, transgenes were cloned into a pShuttle-CMV plasmid (gift from Bert Vogelstein; Addgene plasmid #16403) via Gibson Assembly. PShuttle-CMV plasmids were then digested overnight with MssI (ThermoFisher) and Linearized pShuttle-CMV plasmids were transformed into the final viral backbone using electrocompetent AdEasier-1 cells (gift from Bert Vogelstein; Addgene, #16399). Successful incorporation of pShuttle-CMV construct into AdEasier-1 cells confirmed via digestion with PacI (ThermoFisher). 5000 ng plasmid was then digested at 37℃ overnight, then 85℃ for 10 minutes and transfected in a 3:1 polyethylenimine (PEI, Sigma):DNA ratio into 70% confluent HEK 293A cells (ThermoFisher) in a T-25 flask.

Over the course of 2-4 weeks, fluorescent cells became swollen and budded off the plate. Once approximately 70% of the cells had lifted off the plate, cells were scraped off and spun down at 2000 rpm for 5 minutes in a 15 mL conical tube. The supernatant was aspirated, and cells were resuspended in 1 mL PBS. Cells were then lysed by 3 consecutive quick freeze-thaw cycles in liquid nitrogen, spun down for 5 minutes at 2000 rpm, and supernatant was added to 2qty 70% confluent T-75 flasks. Propagation continued and collection repeated for infection of 10-15cm dishes. After collection and 4 freeze thaw cycles of virus collected from 10-15cm dishes, 8 mL viral supernatant was collected and combined with 4.4 g CsCl (Sigma) in 10 mL PBS. Solution was overlaid with mineral oil and spun at 32,000 rpm at 10℃ for 18 hours. Viral fraction was collected with a syringe and stored in a 1:1 ratio with a storage buffer containing 10 mM Tris, pH 8.0, 100 mM NaCl, 0.1 percent BSA, and 50% glycerol. HUVEC were treated with virus for 16 hours at a 1/1000 final dilution in all cell culture experiments.

### Immunofluorescence and Microscopy

For immunofluorescence imaging, HUVECs were fixed with 4% paraformaldehyde (PFA) for 7 minutes. ECs were then washed three times with PBS and permeabilized with 0.5% Triton-X (Sigma) for 10 minutes. After permeabilization, cells were washed three times with PBS. ECs were then blocked with 2% bovine serum albumin (BSA) for 30 minutes. Once blocked, primary antibodies were incubated for approximately 4-24 hours. Thereafter, primary antibodies were removed, and the cells were washed 3 times with PBS. Secondary antibody with 2% BSA were added and incubated for approximately 1-2 hours, washed 3 times with PBS and mounted on a slide for imaging. For imaging the fibrin-bead assay, first fibroblasts were removed from the clot with a 1-minute trypsin incubation. Following incubation, the trypsin was neutralized with DMEM containing 10% BSA, washed 3 times with PBS, and fixed using 4% PFA for 40 minutes. After fixation, the clot was washed 3 times with PBS, permeabilized with 0.5% Triton-X for 2 hours and then blocked with 2% BSA for 1 hour prior to overnight incubation with primary antibodies. The following day, primary antibodies were removed, and the clot was washed 5 times with PBS and secondary antibody was added with 2% BSA and incubated overnight. Prior to imaging the clot was washed 5 times with PBS. All primary and secondary antibodies are listed in the Supplemental Data. Images were taken on a Nikon Eclipse Ti inverted microscope equipped with a CSU-X1 Yokogawa spinning disk field scanning confocal system and a Hamamatusu EM-CCD digital camera. Cell culture images were captured using a Nikon Plan Apo 60x NA 1.40 oil objective using Olympus type F immersion oil NA 1.518. All images were processed using ImageJ (FIJI).

### Detection of Globular and Filamentous Actin

Globular and filamentous actin ratios were determined by western blot as described by commercially available G-actin/ F-actin *In Vivo* Assay Kit (Cytoskeleton). Globular and filamentous immunocytochemistry was performed as previously described [53]. Briefly, cells were fixed with 4% PFA for 10 minutes and permeabilized in ice cold acetone for 5 minutes and washed. Cells were then incubated for 15 minutes in 2% BSA with globular actin-binding protein GC globulin (Sigma). Following incubation, cells were washed three times in PBS. After washes cells incubated with an anti-GC antibody in BSA for 15 minutes, washed three times, and incubated in anti-rabbit-555 secondary prior to imaging.

### Antibody Feeding Assay

Antibody feeding assay was carried out as previously described [70]. Briefly, cells were moved to 4^0^C for 30 minutes to inhibit endocytosis and then β1-integrin antibody was added to the culture for an additional 30 minutes. Following incubation, cells were washed 3 times with ice cold PBS and moved back into the 37°C degree incubator for 20 minutes. Cells were then fixed with 4% PFA for 8 minutes and washed with PBS. β1-integrin antibody was added once more for 45 minutes to label extracellular integrins, washed 3 times with PBS, and then incubated with the secondary antibody (Alexa 555). The secondary was washed 3 times with PBS and then permeabilized with 0.5% Triton-X for 10 minutes to gain access to the endocytosed β1-integrin pool. Then a secondary antibody (Alexa 488) was added for 20 minutes to label the endocytosed integrins, washed and imaged.

### Wound Healing Assay

Treated cells were moved to Ibidi culture insert plates with a two well silicone insert allowing for a defined cell-free gap. At 3 days post siRNA treatment the silicone insert was removed, and cells were allowed to migrate for 6 hours. Thereafter, cells were fixed, and immunohistochemistry was performed. The distance traveled into the cell free space was measured between groups.

### Immunoblotting & Protein Pull-Down

HUVEC cultures were trypsinized and lysed using Ripa buffer (20 mM Tris-HCl [pH 7.5], 150 mM NaCl, 1 mM Na2EDTA, 1 mM EGTA, 1% NP-40, 1% sodium deoxycholate, 2.5 mM sodium pyrophosphate, 1 mM β-glycerophosphate, 1 mM Na_3_VO_4_, 1 µg/mL leupeptin) containing 1x ProBlock™ Protease Inhibitor Cocktail −50 (GoldBio). Total concentration of protein in lysate was quantified using the Pierce™ BCA Protein Assay Kit measured at 562 nm and compared to a standard curve. 20-50 µg protein was prepared in 0.52 M SDS, 1.2 mM bromothymol blue, 58.6% glycerol, 75 mM Tris pH 6.8, and 0.17 M DTT. Samples were boiled for 10 minutes, then loaded in a 7-12% SDS gel and run at 150 V. Protein was then transferred to Immun-Blot PVDF Membrane (BioRad) at 4℃, 100 V for 1 hour 10 minutes. Blots were blocked in 2% milk proteins for 1 hour, then put in primary antibody at specified concentrations overnight. After 3 10-minute washes with PBS, secondary antibodies at specified concentrations were applied for 4 hours. After 3 additional PBS washes, blots were developed with ProSignal® Pico ECL Spray.

For GGA3 pull-down experiments, GST-GGA3 was grown overnight in 50 mL of Laria-Bertani broth in NiCo21 E Coli (NEB). The following day the overnight culture was transferred to 1L of terrific buffer. The culture was monitored for growth and induced at OD600 with IPTG (GoldBio, 12481) at a final concentration of 100uM. Following induction, bacteria were incubated for an additional 3 hours. Induced cells were collected and pelleted, with the pellet resuspended in cold PBS containing 1mg/ml lysozyme and 1x ProBlock™ Protease Inhibitor Cocktail −50 and then sonicated to lyse bacteria. Cell lysate was clarified by centrifugation and glutathione agarose resin (GoldBio) was added to affinity purify the GST-GGA3. After incubation, agarose resin was washed 2-3 times with PBS and stored at −20°C.

### Zebrafish Transplantation, Microangiography and Gene Editing

Zebrafish transplantations were performed as previously described [71]. Briefly, cells were harvested at the blastula stage from a tg(kdrl:mCherry) line and treated with CRISPR (described below) line using an Eppendorf CellTram and deposited into recipients harboring a tg(kdrl:eGFP) transgene allowing us to distinguish between host and recipient blood vessels.

For microangiography 48 hpf embryos were (anesthetized) with 1% tricaine for approximately 20 minutes prior to perfusion. Embryos were then loaded ventral side up onto an injection agarose facing the injection needle. Qdots (ThermoFisher) were sonicated prior to injection. Qdots were loaded into a pulled capillary needle connected to an Eppendorf CellTram and 1-3μl of perfusion solution was injected into the pericardial cavity. Once successfully perfused, embryos were embedded in 0.7% low melt agarose and imaged promptly. Images were taken on a Nikon Eclipse Ti inverted microscope equipped with a CSU-X1 Yokogawa spinning disk field scanning confocal system and a Hamamatusu EM-CCD digital camera using either Nikon Apo LWD 20x NA 0.95 or Nikon Apo LWD 40x NA 1.15 water objective.

Tol2-mediated transgenesis was used to generate mosaic intersomitic blood vessels as previously described [72, 73]. Briefly, Tol2 transposase mRNA were synthesized (pT3TS-Tol2 was a gift from Stephen Ekker, Addgene plasmid # 31831) [74] using an SP6 RNA polymerase (mMessage Machine, ThermoFisher). A total of 400ng of transposase and 200ng of plasmid vector were combined and brought up to 10μL with phenol red in ddH2O. The mixture was injected into embryos at the 1-2 cell stage. Injected zebrafish were screened for mosaic expression at 48 hpf and imaged.

CRISPR/cas9-mediated knockouts were performed as previously described [54]. Briefly, equal volumes of chemically synthesized AltR® crRNA (100 μM) and tracrRNAr RNA (100 μM) were annealed by heating and gradual cooling to room temperature. Thereafter the 50:50 crRNA:tracrRNA duplex stock solution was further diluted to 25 μM using supplied duplex buffer. Prior to injection 25 μM crRNA:tracrRNA duplex stock solution was mixed with 25 μM Cas9 protein (Alt-R® S.p. Cas9 nuclease, v.3, IDT) stock solution in 20mM HEPES-NaOH (pH 7.5), 350mM KCl, 20% glycerol) and diluted to 5μM by diluting with water. Prior to microinjection, the RNP complex solution was incubated at 37°C, 5 min and then placed on ice. The injection mixture was micro-injected into 1-2 cell stage embryos. Crispant DNA was retrieved via PCR and subjected to sanger sequencing to visualize indel formation.

### Zebrafish Live Imaging and Quantification

All zebrafish presented were imaged at 48hpf. Prior to imaging, embryos were treated with 1% Tricaine for 20 minutes and afterwards embedded in 0.7% low melt agarose. Live imaging of Zebrafish intersomic vessels (ISVs) were performed using the spinning-disk confocal microscopy system mentioned above. ISVs that were analyzed were between the end of the yolk extension and tail. Parameters measured included ISV number, number of non-lumenized vessels (no visible separation between opposing endothelial cells in ISVs), and number of actin accumulations (actin accumulations with a diameter greater than 4μm).

### Scanning Electron Microscopy

Cells fixed for SEM were followed the procedure outlined by Watanabe, et al [75]. Scanning electron microscopy was performed at the University of Colorado Anschutz Medical Campus by Dr. Eric Wortchow.

### Statistical Analysis

Experiments were repeated a minimum of three times. Statistical analysis and graphing were performed using GraphPad Prism. Statistical significance was assessed with a student’s unpaired t-test for a two-group comparison. Multiple group comparisons were carried out using a one-way analysis of variance (ANOVA) followed by a Dunnett multiple comparisons test. Data was scrutinized for normality using Kolmogorov-Smirnov (K-S) test. Zebrafish sex distribution was not adjusted as sex determination did not occur at the stage of development in which the specimens were assayed. Statistical significance set a priori at p<0.05.

## RESULTS

### Rab35 is required for sprouting angiogenesis

To characterize the role of Rab35 in sprouting angiogenesis, we first cloned a fluorescently tagged version of Rab35 into a lentivirus expression system [24]. Thereafter, we transduced ECs and then challenged the cells to sprout in a fibrin-bead assay [25, 26]. Fibrin-bead sprouts demonstrate excellent angiogenic characteristics, reproducing the most salient sprouting behaviors, such as branching, lumenogenesis, anastomosis and tip/stalk cell signaling (**Fig. 1A**) [7, 8, 27]. Rab35 in 3-dimensional (3D) sprouts demonstrated strong membrane localization, co-localizing with apical marker podocalyxin and luminal actin, opposite basally located β1-integrin (**Fig. 1B, Movie 1**). To test whether Rab35 was necessary for endothelial sprouting, we knocked down Rab35 using siRNA (**Fig. 1C**). Loss of Rab35 reduced sprout length and sprouts per bead by ~50%, with a significant increase in the percentage of non-lumenized sprouts (**Fig. 1D-G**). Morphologically, the sprouts appeared stubby, non-lumenized and generally dysmorphic compared with controls (**Fig. 1D**). These results indicate that Rab35 is required for proper sprout development.

**Figure 1.**
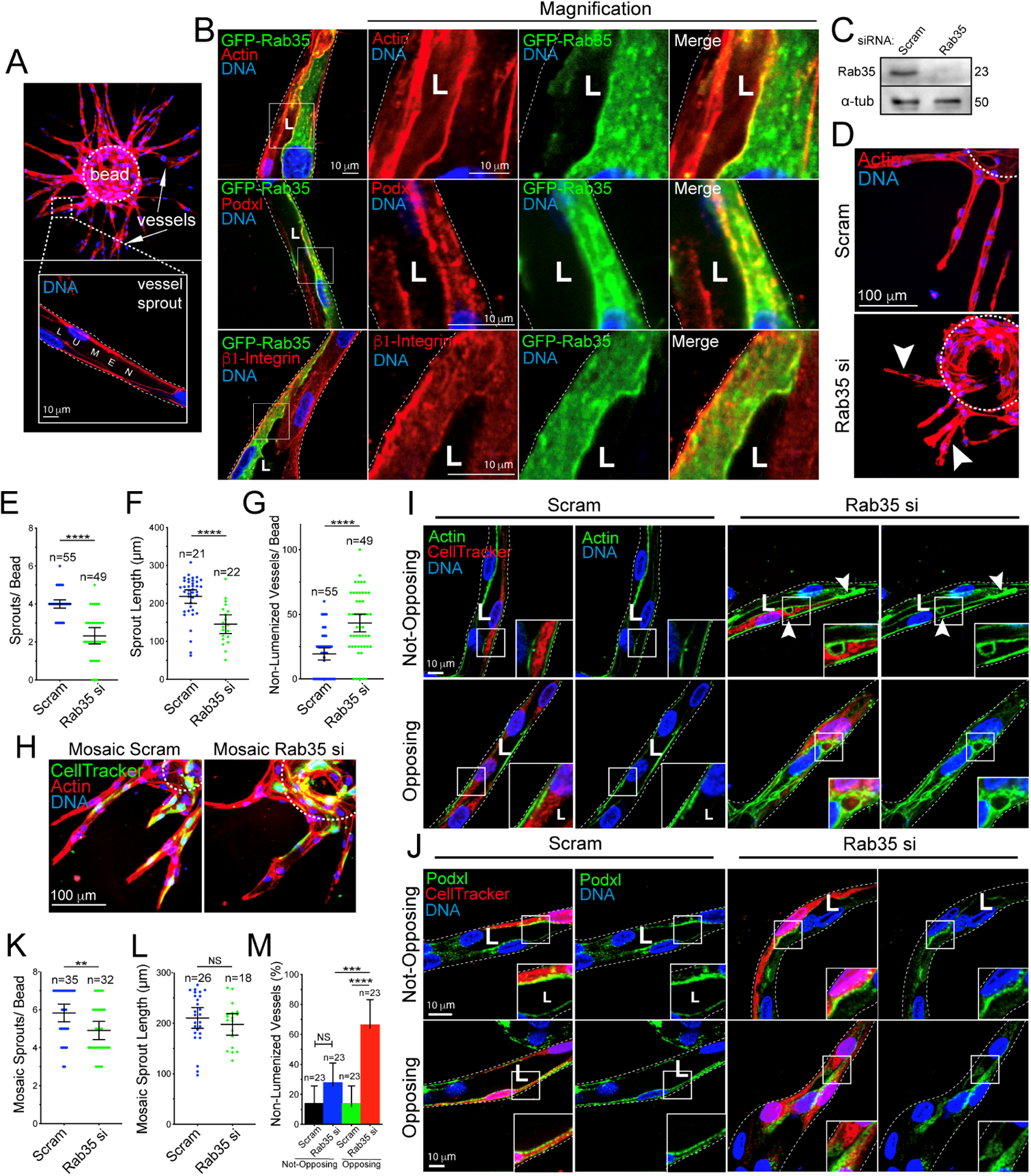
Rab35 is an apical membrane protein required for sprout formation. (**A**) Representative images of the fibrin-bead assay (FBA) at low and high magnification. Arrows mark sprout structures. Inset depicts lumenized sprout. (**B**) GFP-Rab35 localization in endothelial sprouts with actin (top panels), podocalyxin (Podxl, middle panels), and β1-integrin (bottom panels). (**C**) Western blot confirmation of siRNA knockdown of Rab35. (**D**) Representative image of scramble (Scram) control and Rab35 siRNA (si) knock-down (KD) sprouts. Arrowheads denote short and non-lumenized sprouts. Dashed lines outline the microbead. (**E-G**) Graphs of indicated sprouting parameters between groups. (**H**) Representative images of sprout morphology of mosaic Scram and Rab35 KD cells, green indicates cell tracker of siRNA treated cells. (**I,J**) Representative images of non-opposing (top panels, an isolated siRNA treated cell) and opposing (bottom panels, two adjacent siRNA treated cells) cells stained as indicated. Arrowheads denote aberrant actin accumulations (**K-M**) Quantification of indicated parameters across groups. In all images L denotes lumen. ** p<0.01, *** p<0.001, **** p < 0.0001, NS=Non-Significant. Error bars represent 95% confidence intervals. N=number of sprouts. Insets are areas of higher magnification. White dotted lines mark sprout exterior. All experiments were done using Human umbilical vein endothelial cells in triplicate.

Given Rab35 depletion exhibited such a profound impact on sprouting parameters, we stained for various cytoskeletal, apical and basal markers to determine if Rab35 was affecting specific polarity pathways or producing a more global cellular defect. Imaging for VE-cadherin (cell-cell junctions), podocalyxin, β1-integrin (basal membrane), moesin (cytoskeletal, apical membrane), synaptotagmin-like protein-2 (apical membrane) and phosphorylated-Tie2 (apical membrane) revealed that Rab35 knockdown affected all protein localization (**Fig. S1A**), suggesting that loss of Rab35 globally disturbs cell polarity programs. Emblematic of this was the significant lack of lumen formation and the increase in discontinuous vacuoles in the Rab35 depleted condition (**Fig. S1B**), as lumenogenesis requires proper apicobasal signaling to form [7, 28]. We also observed that Rab35 knockdown reduced the number of nuclei per sprout, indicating the presence of cell division defects in line with other reports [16, 29–31] (**Fig. S1C**). Overall, this data suggests that Rab35 plays a significant role in establishing cell polarity during angiogenic sprouting.

We next employed a mosaic approach to determine the cell autonomous nature of Rab35 depletion in a sprout collective. To do so, we treated ECs with either Rab35 siRNA or a scramble control. Thereafter, the knockdown population was marked with cell-tracker and mixed 50:50 with wild-type ECs. The resulting mosaic sprouts contained a mixture of siRNA-treated and untreated ECs (**Fig. 1H**). Cells contained within sprouts were then binned into two categories: 1) not-opposing, an isolated siRNA-treated cell; or 2) opposing, two adjacent siRNA-treated ECs (**Fig. 1I,J**). Our results demonstrate that Rab35 knockdown in not-opposing ECs contained actin-labeled vacuolations and polarity defects as indicated by a reduction in lumen formation compared with scramble-treated controls (**Fig. 1I-M**). For Rab35 depleted ECs in the opposing orientation defects were even more pronounced with complete lumen failures at these sites, while also exhibiting multiple vacuolations and polarity defects (**Fig. 1I-M**). Overall, these results indicate that Rab35 is cell autonomous and is required for EC polarity.

### Rab35 resides at the apical membrane during sprouting

As the loss of Rab35 produced such a profound effect on EC sprouting, we sought to better understand its cellular localization to gain insight into its potential function. In sprouts, quantification of Rab35 enrichment between different cellular compartments showed a preference for the apical membrane for wild-type (WT) and constitutively active (CA) Rab35 variants, while the dominant-negative (DN) Rab35 mutant resided in the cytoplasm (**Fig. 2A**). In this regard, subcellular imaging of WT and CA Rab35 showed a strong colocalization with apical podocalyxin (**Fig. 2B**). Similar to loss of Rab35, expression of the DN Rab35 also produced polarity defects, such as mislocalization of podocalyxin and large actin accumulations (**Fig. 2B**). To more conclusively assign Rab35 phenotypes, we performed several rescue assays by knocking down the endogenous Rab35 and then over-expressing Rab35 variants in sprouts. Expression of WT or CA Rab35 decreased the number of non-lumenized sites in sprouts compared to endogenous Rab35 knockdown alone expressing a GFP control, but not to levels in the scramble treated group (**Fig. 2C,D; S2A,B**). Rab35 knockdown and expression of the DN Rab35 mutant showed the highest increase in dysmorphic sprouts, exhibiting numerous accumulations of actin puncta and lumen defects, again suggesting Rab35 is necessary for sprout function.

**Figure 2.**
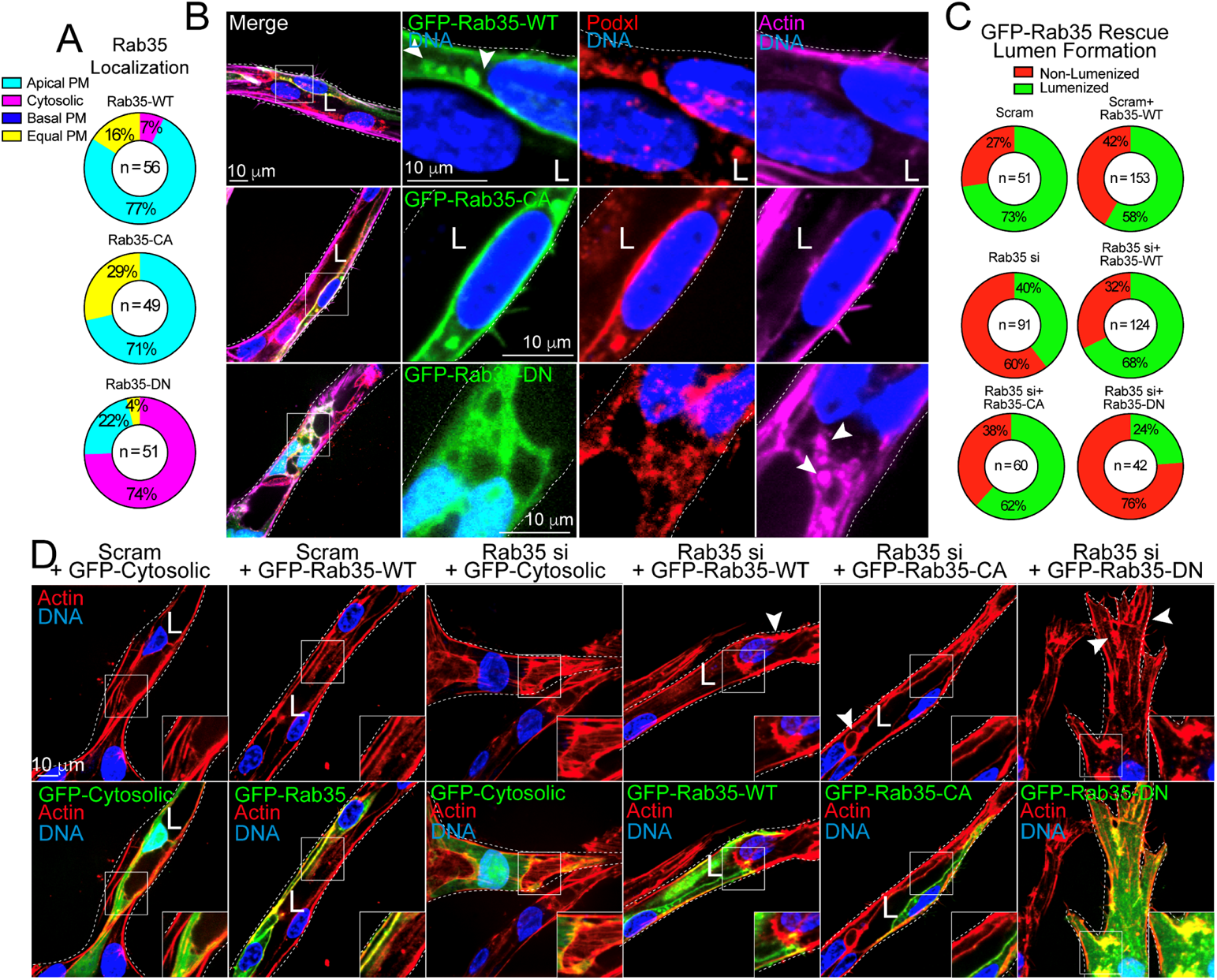
Rab35 mutant localization and rescue in endothelial sprouts. (**A**) Quantification of GFP-Rab35 wild-type (WT), constitutively active (CA), and dominant-negative (DN) localization in endothelial sprouts. Apical plasma membrane (PM, uniformly localized to apical membrane), basal PM (Rab35 uniformly located at the basal membrane), cytosolic (localized in the cytoplasm), equal PM (Rab35 equally distributed between the apical and basal membranes). N= number of cells. (**B**) GFP-Rab35 WT (top panels), CA (middle panels), and DN (bottom panels) localization in endothelial sprouts. Co-staining with Podocalyxin (Podxl) and actin. Arrowheads in top panels denote Rab35 apical localization and puncta. Arrowheads in bottom panels denote abnormal accumulations of actin. (**C**) Quantification of lumen formation in described conditions. N=number of sprouts. (**D**) Representative images of Rab35 KD sprouts rescued with either GFP-cytosolic, (control), or GFP-Rab35-WT/CA/DN. Arrowheads denote actin accumulations. White dotted lines mark sprout exterior. L denotes lumen in all images. Insets are areas of higher magnification. All experiments were done using Human umbilical vein endothelial cells in triplicate.

Within the sprout body, Rab35 also localized to actin at cytokinetic bridges as previously described [16, 29, 31], but had no preference for filopodia extensions or tip-cell positioning (**Fig. S3A,B**). In 2D culture, we also observed that Rab35 modestly colocalized with filamentous actin in a monolayer; however, this association was reduced in migratory cells (**Fig. S3C,D**). Previous reports have implicated Rab35 in Wiebel Palade Body (WPB) granule release [32]. Although loss of Rab35 may alter WPB secretion, likely due to the impact on cell polarity, in our hands Rab35 did not colocalize with these structures in 2D or 3D culture systems (**Fig. S3E,F**). These results indicate that Rab35 is largely localized to the apical membrane in its active form as well as areas of high actin density.

Previous literature in epithelial tissue has reported that Rab35 participates in trafficking of podocalyxin to the apical membrane [15, 16]. In the sprouting model, we observed a strong colocalization of Rab35 and podocalyxin at the apical membrane as well as mislocalization of podocalyxin in the absence of Rab35. This data could be interpreted as a loss of, or defective, podocalyxin trafficking given Rab35’s previous association with this pathway. As colocalization of podocalyxin and Rab35 at the apical membrane could be circumstantial as many proteins localize to the apical membrane during lumenogenesis, we overexpressed TagRFP-Rab35 and stained for endogenous podocalyxin in 2D culture and did not detect any significant signal overlap (**Fig. S4A**). Previous literature showed that Rab35 directly binds to the cytoplasmic tail of podocalyxin [16]. Overexpression of the human podocalyxin cytoplasmic domain (residues 476–551) and TagRFP-Rab35 also did not show any obvious association (**Fig. S4A**). To further probe for this previously reported binding between Rab35 and podocalyxin, we engineered a mitochondrial-targeted Rab35 to test what proteins or complexes bind Rab35 and are then ‘pulled’ along to mitochondria. Expression of WT or CA mitochondrial-targeted Rab35 did not show any association with endogenous podocalyxin or overexpression of its cytoplasmic tail domain (**Fig. S4B,C**). We next reasoned if mistrafficking of podocalyxin by way of Rab35 depletion was the predominant mechanism underpinning the sprouting defects, then knocking down podocalyxin would produce a similar phenotype as compared with loss of Rab35. Knockdown of podocalyxin did not phenocopy Rab35-mediated sprouting defects (**Fig. S4D-I**). The only exception was that podocalyxin knockdown increased the percentage of non-lumenized sprouts compared with controls. Overall, our data suggests that Rab35 does not directly participate in podocalyxin trafficking in ECs; however, loss of Rab35 distorts podocalyxin’s localization to the apical membrane likely due to other alterations in cell polarity.

### Rab35 interacts with ACAP2 in endothelial cells

To take a more holistic approach in determining how Rab35 functions in endothelial tissue, we performed a functional screen by knocking down the most highly cited Rab35 effectors singly and in combination, to determine if any effector combination phenocopied Rab35 sprouting defects (**Fig. 3A,B**) [15, 17–19, 30, 32–35]. First, we found that Rab35 itself did not produced a significant effect on 2D cell motility, suggesting the primary defect in sprouting may be due to altered apicobasal polarity only detectable in the 3D sprout environment (**Fig. S5A,C**). As Rab35 and ACAP2 have been shown to affect the integrin recycling pathway via their association with Arf6, we also assayed for integrin recycling as defective integrin signaling could also affect cell polarity. As compared with the scramble controls, knockdown of Rab35 and OCRL significantly increased integrin recycling, while ACAP2 and MICAL-L1 had no effect (**Fig. S5B,D**).

**Figure 3.**
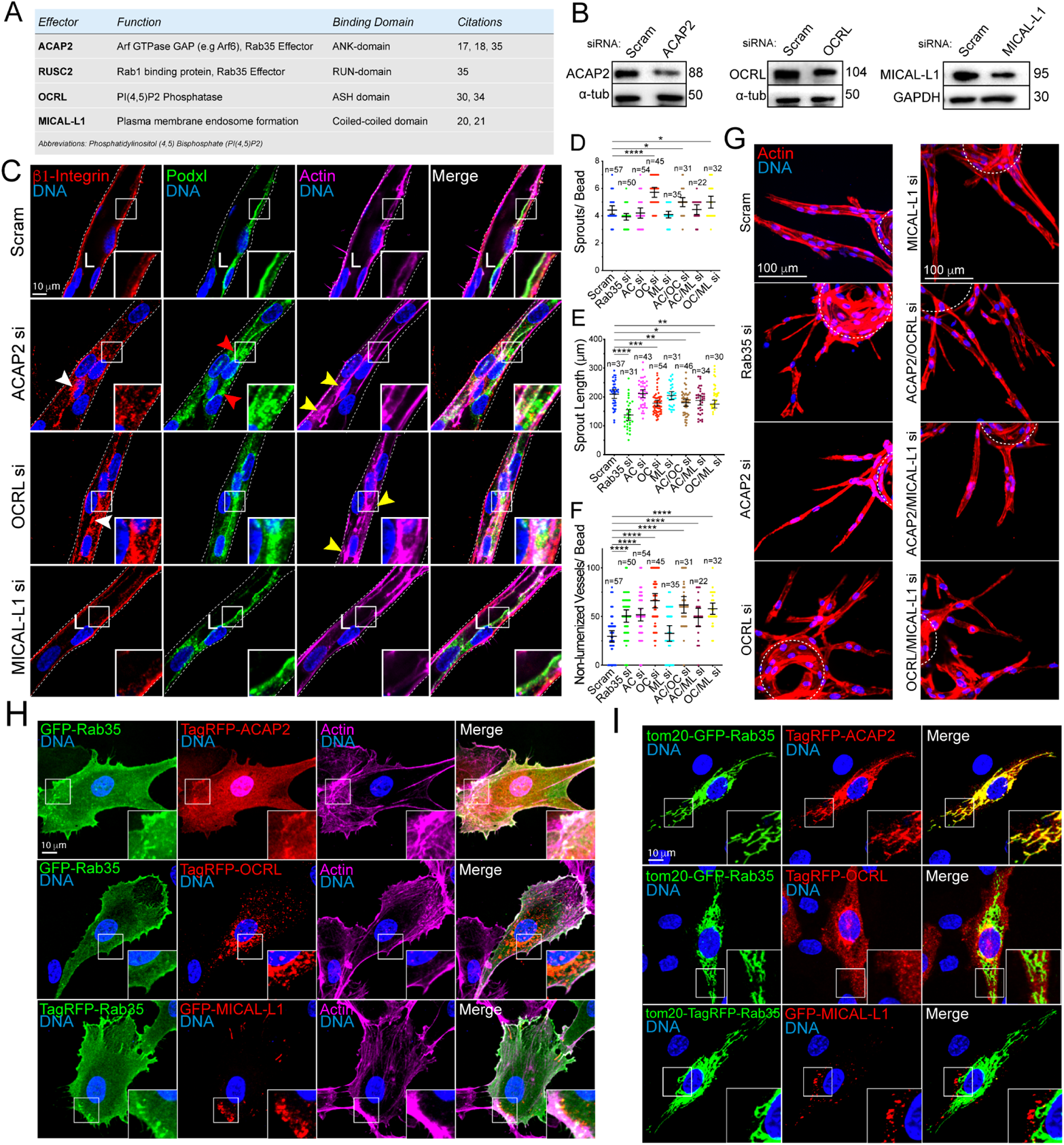
Rab35 effector localization and necessity to endothelial sprout formation. (**A**) Table listing each effector, respective function, and citations. (**B**) ACAP2, OCRL, and MICAL-L1 knockdown (KD) validation by western blotting. (**C**) Representative images of siRNA (si)-mediated KD of each effector. White arrowhead denotes abnormal localization of β1-integrin. Red arrowheads denote abnormal podocalyxin (Podxl) localization. Yellow arrowheads denote abnormal actin accumulations. White dotted lines mark sprout exterior. (**D-F**) Graphs of indicated sprout parameters between groups. ACAP2 (AC), OCRL (OC), and MICAL-L1 (ML). N= number of sprouts. (**G**) Representative images of sprout morphology between indicated groups. Dashed lines outline microbeads. (**H**) Two-dimensional localization of GFP-Rab35 or TagRFP-Rab35 with indicated effectors and stained for actin. (**I**) Representative images of mitochondrial mis-localization experiment. Rab35 was unnaturally tethered to the mitochondria with a tom20 N-terminal tag to test if indicated effectors were also mislocalized to the mitochondria. In all images L denotes lumen. * p < 0.05, ** p<0.01, *** p<0.001, **** p < 0.0001, NS=Non-Significant. Error bars represent 95% confidence intervals. N=number of sprouts. Insets are areas of higher magnification. All experiments were done using Human umbilical vein endothelial cells in triplicate.

Next, we determined that RUSC protein levels were not detectable in ECs, thus was excluded from our screen (**Fig. S6F**). ACAP2, OCRL, or MICAL-L1 or any combination of knockdown targeting these proteins, demonstrated the greatest phenotypic similarity to Rab35 knockdown with regard to sprouting parameters (**Fig. 3C-G**). Upon closer inspection, both ACAP2 and OCRL knockdowns were associated with elevated frequencies of non-lumenized sprouts with disorganized actin, although to a lesser extent than compared with Rab35 (**Fig. 3C**). These results suggest that ACAP2 and OCRL potentially resemble a Rab35 sprouting defect.

Both ACAP2 and OCRL have been reported to directly bind Rab35 [17–19, 30, 32, 35]; however, this interaction has not been validated in ECs. First, we overexpressed tagged versions of ACAP2, OCRL and MICAL-L1 to visualize their localization patterns with Rab35 in ECs. Rab35 and ACAP2 strongly colocalized to the plasma membrane, while Rab35 did not show strong localization with ORCL or MICAL-L1 (**Fig. 3H**). Further testing for potential interactions, we again used the mitochondrial-targeted Rab35 to visualize any physical association between Rab35 and these previously published effectors. Co-expression of WT and CA Tom20-Rab35 with ACAP2 demonstrated strong colocalization at the mitochondria, while the DN Rab35 showed no significant binding of ACAP2 (**Fig. 3I; S6A,B**). We performed this same experiment using ACAP2 with the ankyrin repeat domain deleted and observed no binding, indicating Rab35 directly interacts with this domain (**Fig. S6B**). As a control, we also co-expressed a tom20-Rab27a and ACAP2 and observed no mislocalization of ACAP2 (**Fig. S6C**), suggesting ACAP2’s affinity for Rab35 is specific. Co-expression of WT, CA or DN Tom20-Rab35 with OCRL or MICAL-L1 did not show any colocalization at the mitochondria, signifying a lack of binding (**Fig. 3I; S6D,E**). These results demonstrate that ACAP2, not OCRL or MICAL-L1, directly interacts with Rab35 in endothelial tissue.

### Rab35 does not impact Arf6 activity in endothelial cells

Previous literature has shown that ACAP2 works as a GTPase activating protein (GAP) with Rab35 to inactivate the GTPase Arf6 [17, 18, 33]. Arf6 has been shown to be involved with actin remodeling and integrin recycling [19, 36–38]. To test if this association exists in ECs, we first determined the localization of Arf6 relative to Rab35 and ACAP2 in 2D culture. Cells expressing tagged Rab35 and Arf6, or ACAP2 and Arf6 demonstrated modest colocalization throughout the cell with the greatest colocalization at the cell cortex (**Fig. S7A**). Using WT, CA and DN versions of Arf6, we stained for actin to determine if Arf6, like Rab35, associated with actin structures. Similar to Rab35, in peripheral membrane protrusions, WT and CA Arf6 demonstrated moderate colocalization with actin; however, this association did not persist on filamentous actin located towards the cell interior (**Fig. S7B**). In sprouts, Arf6 showed weak localization to the apical membrane as compared to Rab35 (**Fig. S7C**). Once again using mitochondrial-mistargeting, we tested for binding between Arf6 and Rab35. Mitochondrial-targeted Rab35 did not pull Arf6, indicating a lack of binding interaction (**Fig. S7D**). To further confirm this, we used the Tom20 epitope to target ACAP2 to the mitochondria to determine if Arf6 interacts with ACAP2. Our results show that Arf6 does not localize to the mitochondria, indicating ACAP2 does not strongly interact with Arf6 in ECs (**Fig. S7D**). We reasoned that the lack of binding between Arf6 and ACAP2 could be due to an insufficiency of Rab35, as Rab35 is hypothesized to regulate ACAP2’s availability to act upon Arf6 [17]. Therefore, we simultaneously expressed Tom20-Rab35, TagRFP-ACAP2 and HA-Arf6 in hopes that the Rab35 bound to ACAP2 would recruit Arf6 to the mitochondria. Our results demonstrate that Arf6 did not localize with mitochondrial Rab35 and ACAP2, suggesting ACAP2 does not directly act upon Arf6, or that this signaling does not require a robust binding interaction in ECs (**Fig. S7E**).

Due to the wealth of literature demonstrating loss of Rab35 increases Arf6 activity in non-endothelial tissues, we sought to confirm this signaling interaction biochemically. To do so, we first expressed WT, CA, and DN versions of Arf6 in ECs and used recombinant GGA3 to pulldown the active form of Arf6 as others have reported [39]. Pulldown using GGA3 demonstrated more binding with the CA mutant as compared with the WT and DN versions of Arf6, validating this approach for testing Arf6 activity (**Fig. S7F**). Next, we knocked down and over-expressed Rab35 in ECs and then probed for active Arf6. Knockdown of Rab35 or overexpression of Rab35 did not significantly alter Arf6 activity in ECs (**Fig. S7G**). These results would suggest that loss of Rab35 does not affect Arf6 activation. Previous literature reported that knockdown of Rab35 promoted Arf6 activity; thus, we next tested if overactivation of Arf6 would phenocopy the Rab35 loss of function sprouting phenotype to more thoroughly factor out this signaling pathway. Moving to Arf6 overexpression in sprouts, we observed that both WT and CA Arf6 marginally affected sprouting parameters with the WT and CA Arf6 increasing the frequency of lumen failures compared with the DN version (**Fig. S7H)**. A primary phenotype in sprouts deficient in Rab35 was abundant actin aggregates and the presence of non-apical podocalyxin. Inconsistent with these observations, ECs expressing CA Arf6 demonstrated normal actin architecture as well as typical podocalyxin apical deposition (**Fig. S7I**). These results suggest that overactivation of Arf6 due to loss of Rab35 is likely not the causative pathway promoting sprouting defects.

### DENNd1c is required for Rab35 function

We were intrigued by the idea that other roles of Rab35 were being unaccounted for as Arf6 activation was largely unaffected by loss of Rab35 in ECs. Earlier we observed that Rab35 colocalized with actin. Additionally, a consistent phenotype we observed was impaired actin organization, marked by actin aggregates in Rab35 knockdown sprouts. To this end, Rab35 has 3 GEFs, DENNd1a-c [40–43]. DENNd1c has been shown to play a more uncharacteristic role, being less involved with GTP hydrolysis, but demonstrating the lone ability to bind to both globular and filamentous actin, mediating Rab35 localization to these microfilaments [41]. Exploring this association, we knocked down DENNd1a-c individually and in combination. Loss of DENNd1a and DENNd1b did not produce any significant impact on sprouting morphology; however, knockdown of DENNd1c alone resulted in growth of dysmorphic sprouts mirroring Rab35 loss of function (**Fig. 4A-E; Movie 2,3**). Knockdown of all DENNd1s produced the greatest effect on sprouting behaviors, presumably because the GEF activity provided by DENNd1a/b was also lost (**Fig. 4D-F**). We also confirmed that knocking down any given DENNd1 did not result in a compensatory increase in expression of the remaining DENNd1s (**Fig. S8B**). Staining for actin demonstrated that DENNd1c knockdown produced the greatest number of aberrant actin accumulations similar to the Rab35 knockdown phenotype (**Fig. 4C,G**). We next cloned and tagged DENNd1c to visualize its cellular localization with Rab35. DENNd1c and Rab35 showed strong colocalization on actin in 2D cell culture (**Fig. S8A**). We also expressed Rab35 with the integral actin protein Arp2 that mediates actin filament branching [44], Rab35 localized to areas of active polymerization marked by Arp2/Rab35 localization. Rab35, Arp2 and filamentous actin colocalized in many areas (**Fig. S8A**). To explore if DENNd1c, per se, was responsible for tethering Rab35 to actin, we individually knocked down all three DENNd1s and quantified the relative amount of Rab35 uniformly localized at the plasma membrane, accumulated at the plasma membrane or in the cytoplasm. DENNd1c knockdown exhibited the greatest increase in apical plasma membrane accumulations compared with DENNd1a or DENNd1b (**Fig. 4H**). These data indicate that loss of DENNd1c phenocopies the Rab35 knockdown effect on sprouting parameters and the actin cytoskeleton.

**Figure 4.**
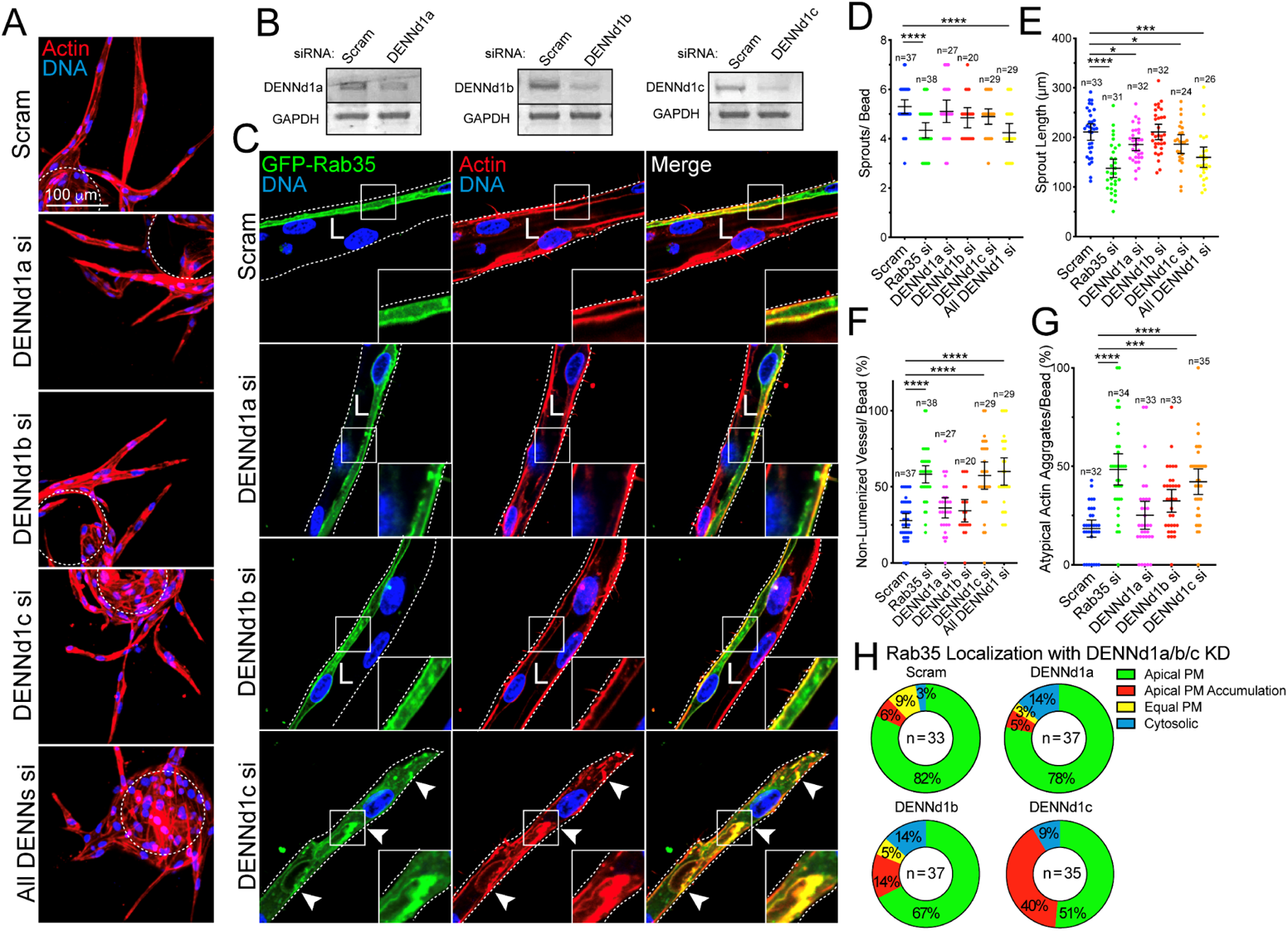
DENNd1c is required for sprouting and Rab35 function. (**A**) Sprout morphology of scramble (Scram), DENNd1a-c and combined siRNA (si)-treated sprouts, stained with actin to denote the general morphology. Dashed line denotes microbead. (**B**) Knockdown confirmations for DENNd1a-c by RT-PCR. (**C**) Representative images of siRNA knockdowns described in A with GFP-Rab35 localization. L denotes lumen and arrowheads denote abnormal actin accumulations. White dotted lines mark sprout exterior. (**D-G**) Graphs of indicated sprout parameters across groups. N=number of sprouts. (**H**) GFP-Rab35 localization in DENNd1a-c siRNA-treated sprouts. Localizations were binned to apical plasma membrane (PM, Rab35 >80% at apical membrane), apical PM accumulations (non-continuous, visible puncta), equal PM (equally enriched at apical and basal membranes), and cytosolic. N=number of sprouts. * p < 0.05, *** p<0.001, **** p < 0.0001, NS=Non-Significant. Error bars represent 95% confidence intervals. Insets are areas of higher magnification. All experiments were done using Human umbilical vein endothelial cells in triplicate.

### Rab35 and DENNd1c localize to sites of actin polymerization

We next sought to comprehensively characterize the association between Rab35, DENNd1c and branched actin located at the cell periphery. To do so, we again overexpressed the actin-specific protein Arp2 as a marker of active actin polymerization [45]. Both Rab35 and DENNd1c demonstrated strong colocalization to Arp2 and the underlying actin **(Fig. 5A**). To specifically perturb the branched actin network, we next treated cells with the Arp2/3 inhibitor CK-666 [46] and then determined the effect on Rab35 and DENNd1c localization. In 3D sprouts, inhibition of branching actin resulted in accumulations of actin similar to the Rab35 knockdown phenotype (**Fig. 5B, S8D; Movie 4,5**). In 2D culture, CK-666 treatment rapidly depleted actin at the cell cortex (**Fig. 5C**). Rab35 prior to CK-666 administration exhibited a uniform distribution in the plasma membrane with enrichment at sites of actin accumulation adjacent to the cell periphery. However, after inhibition of branched actin formation Rab35 collapsed into discrete puncta scattered throughout the cytoplasm. Interestingly, CK-666 treatment created large, presumably globular, actin vacuoles which were then surrounded by Rab35 (**Fig. 5C, S8E; Movie 6**); we believe these structures are analogous to the actin accumulations observed in sprouts when Rab35 is depleted. As a control we performed the same experiment with Rab11a and did not observe any alteration in Rab11a localization with CK-666 treatment (**Fig. S8C**), suggesting not all Rabs are dependent on actin for their localization. Using the same approach with DENNd1c, we observed, again, DENNd1c was highly enriched at cortical actin; however, treatment with CK-666 effectively depleted DENNd1c from this actin population (**Fig. 5D**). Unlike Rab35, CK-666 treatment did not cause the formation of puncta, but the redistribution of DENNd1c to unaffected actin, such as filamentous actin towards the cell interior (**Fig. 5D; Movie 7**). As a control, we treated cells with CK-666 expressing both Rab35 and Arp2. As expected, Arp2 was no longer located on actin, collapsing into puncta, while remaining adjacent to Rab35 (**Fig. 5E; Movie 8**). These data suggest that Rab35 and DENNd1c are recruited to polymerizing actin filaments. Additionally, in the absence of active actin polymerization Rab35 collapses into vesicular structures.

**Figure 5.**
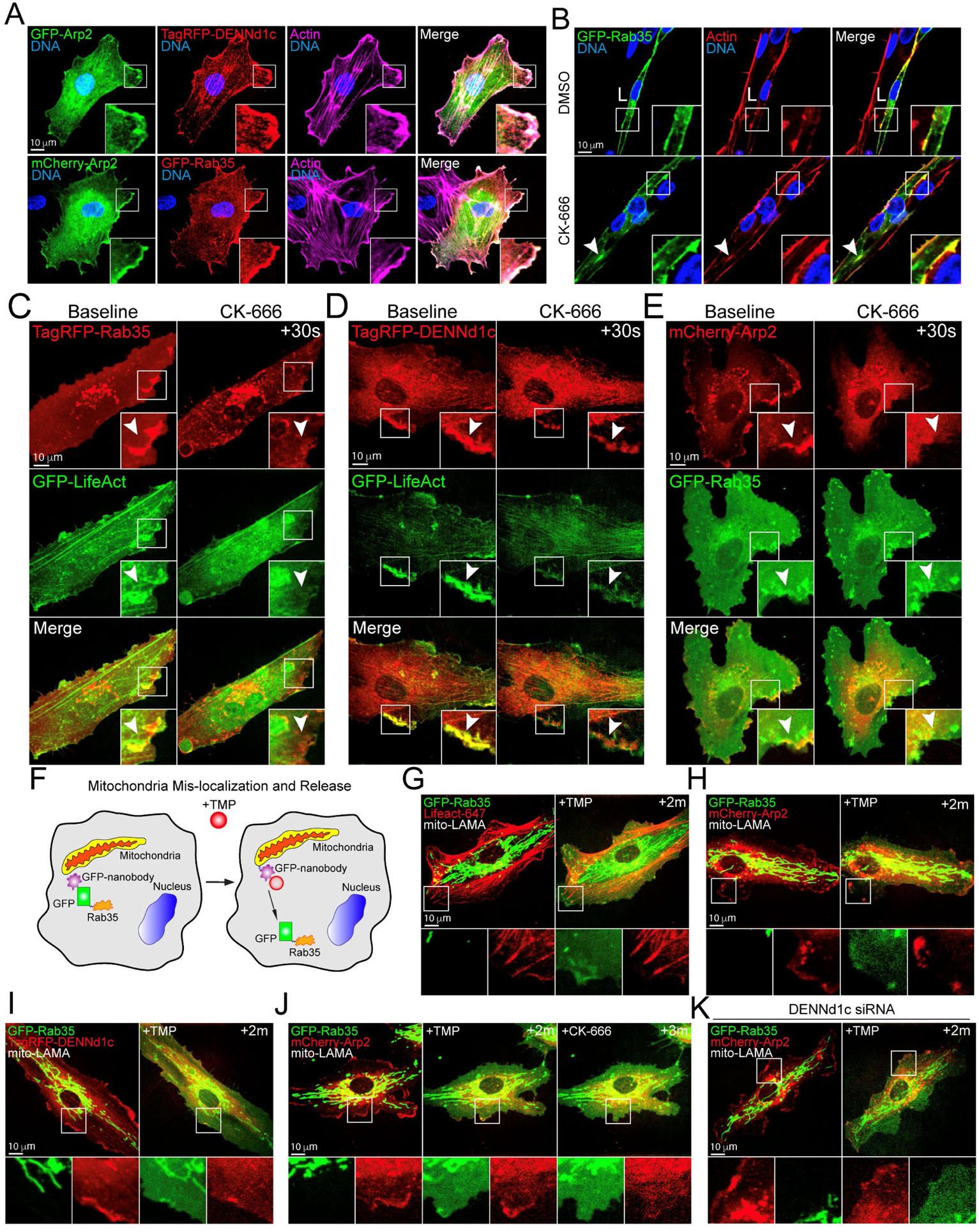
Rab35 localizes to cortical actin. (**A**) Two-dimensional localization of GFP-Arp2 with DENNd1c (top panels) and GFP-Rab35 (bottom panels). (**B**) Representative images of DMSO and CK-666 (Arp Inhibitor) treated sprouts expressing GFP-Rab35. L denotes lumen. (**C,D**) Live imaging of GFP-Rab35 or tagRFP-DENNd1c with TagRFP647-LifeAct at baseline and after treatment with CK-666. White arrowheads denote disappearance of Rab35 puncta over time. (**E**) Representative live-images of a cell expressing mCherry-Arp2 and GFP-Rab35 before and after CK-666 treatment White arrowheads denote disappearance of Rab35 puncta over time. (**F**) Cartoon of a mitochondria-localized GFP-nanobody and controlled release of GFP-Rab35 upon treatment with Trimethoprim (TMP). In the absence of TMP the nanobody sequesters GFP or GFP-tagged proteins. In the presence of TMP the GFP cargo is released. (**G**) Live-image of a cell expressing GFP-Rab35, TagRFP647 (647)-LifeAct and ligand-modulated antibody fragments targeted to the mitochondria (mito-LAMA) before and after TMP administration. (**H**) Live-image of a cell expressing GFP-Rab35, mCherry-Arp2 and mito-LAMA before and after TMP administration. (**I**) Live-image of a cell expressing GFP-Rab35, TagRFP-DENNd1c and mito-LAMA before and after TMP administration. (**J**) Live-image of a cell expressing GFP-Rab35, mCherry-Arp2 and mito-LAMA before and after TMP administration and then treated with CK-666. (**K**) Live-image of a cell expressing GFP-Rab35, mCherry-Arp2 and mito-LAMA treated with DENNd1c siRNA (si) before and after TMP administration. Insets are areas of higher magnification. All experiments were done using Human umbilical vein endothelial cells in triplicate.

To visualize Rab35’s temporospatial recruitment to cortical actin, we employed a chemically switchable GFP-binding nanobody, termed ligand-modulated antibody fragments (LAMAs) [47]. This method allowed us to sequester GFP-tagged Rab35 at the mitochondria and then rapidly release the protein upon drug treatment, enabling dynamic imaging of localization patterns (**Fig. 5F**). Using GFP-Rab35, LAMA and TagRFP647-LifeAct [48] expressing cells, we release GFP-Rab35 from mitochondria and live-imaged its localization preferences. Our data shows that Rab35 quickly localizes to the cell periphery following trimethoprim (TMP) treatment. Of note, Rab35 did not co-localize with longer-lived filamentous actin, as the LifeAct probe primarily decorates this population (**Fig. 5G, Movie 9**). When repeated with tagged Arp2, Rab35 quickly (~2min) localized to Arp2 puncta on the cell cortex (**Fig. 5H; Movie 10**). Rab35 also demonstrated a preference for sites of DENNd1c when released from the mitochondria (**Fig. 5I, Movie 11**). Next, we released Rab35 and imaged its localization to Arp2, and then immediately treated with CK-666 to determine how this association would be affected. Administration of CK-666 rapidly dissociated Rab35 and Arp2 at the cortex (**Fig. 5J; Movie 12**). Lastly, to test if DENNd1c was responsible for recruiting Rab35 to branched actin, we knocked down DENNd1c and repeated the LAMA localization experiments. Upon release, Rab35 showed a reduction in its ability to localize to cortical Arp2, suggesting DENNd1c is important for this interaction (**Fig. 5K**; **Movie 13**). Overall, these data suggest that Rab35 is rapidly recruited to the cortex and is anchored to actin filaments by DENNd1c.

### Rab35 promotes actin assembly

Our previous data indicates that in the absence of Rab35 global apicobasal polarity is affected, which is likely due to significant alterations in the actin cytoskeleton. Also, Rab35 colocalized with actin and actin polymerizing protein Arp2. Thus, our next aim was to test whether Rab35 affected actin polymerization, per se. Prior literature indicates that Rab35 would increase actin polymerization via its purported trafficking interactions with Cdc42 and Rac1 [40, 41, 43]; however, others have claimed Rab35 may act as a brake for actin polymerization through its association with MICAL-L1 in non-endothelial tissues [21].To begin to explore how Rab35 impacts actin in ECs, we transfected Rab35 variants WT, CA and DN into freely migrating ECs. It is well-established that lamellipodia protrusions (membrane movement away from the cell body) and retractions (membrane movement towards the cell interior) are primarily mediated by local actin assembly and disassembly [49, 50]. Attempting to monitor global lamellipodia dynamics in an unbiased fashion, we employed the open source software ADAPT [51]. Our analysis determined that only the Rab35-CA mutant significantly increased both the cells protrusive and retractive capabilities, a finding in line with enhanced migration (**Fig. 6A,B**). Interestingly, knockdown of Rab35 did not shift membrane dynamics significantly, potentially suggesting Rab35-based actin regulation may play a more critical role in 3D sprouting.

**Figure 6.**
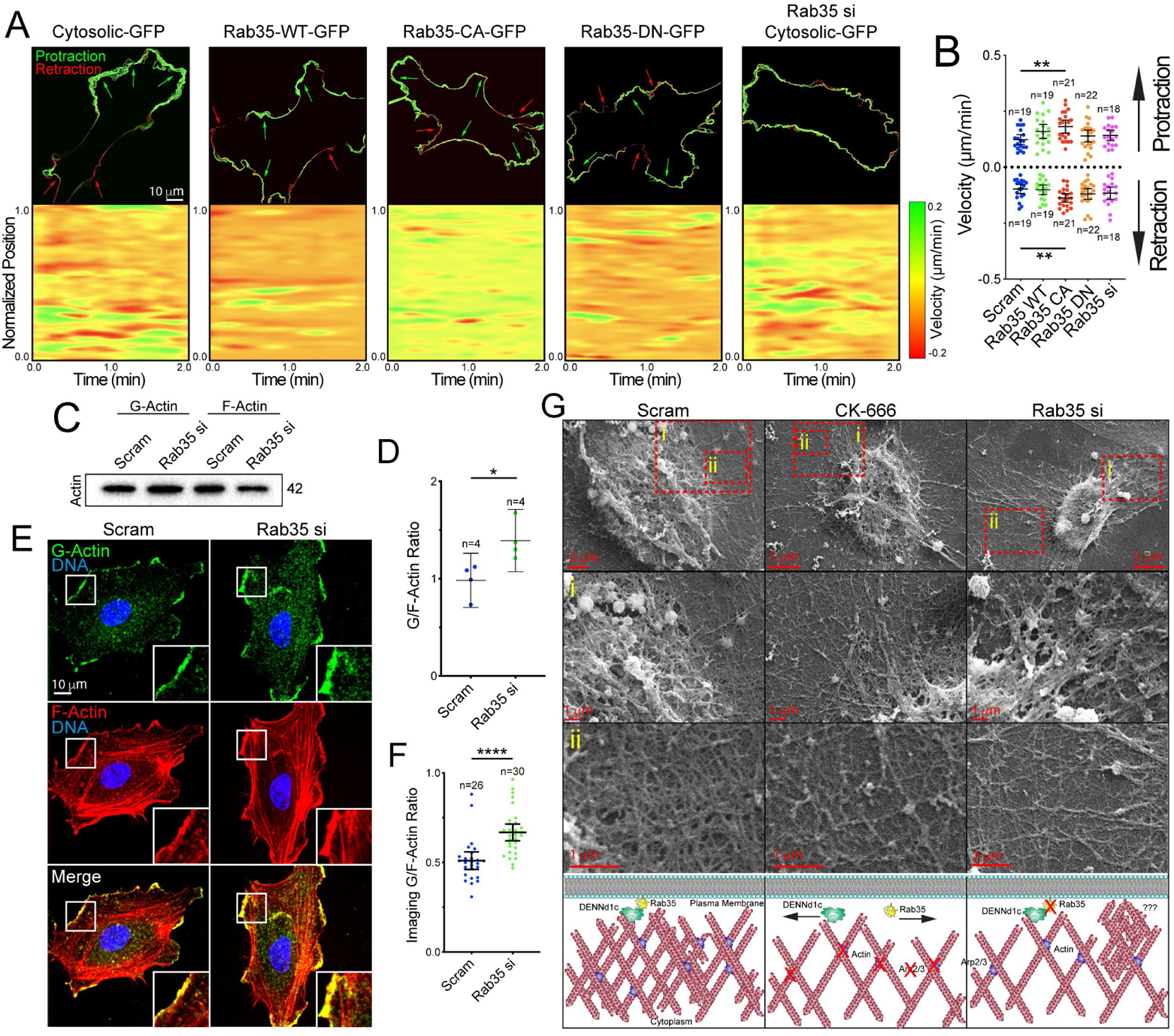
Rab35 regulates actin dynamics. (**A**) Top panels depict change in membrane velocities over time in described conditions. Green represents protraction and red represents retraction of the membrane. Arrows indicate directionality. The bottom panels are heat maps in which the Red is indicative of retractive movement and green is protractive movement over time. Yellow indicates no change in velocity. (**B**) Quantification of cell membrane velocities between indicated groups. Above the dashed line is the protractive velocities and below the dashed line is retractive velocities. N=number of cells. (**C**) Western blot of globular and filamentous actin in siRNA (si)-treated groups. (**D**) Quantification of the ratio of globular to filamentous actin from blots represented in panel C. (**E**) Representative images of cells stained for globular and filamentous actin between indicated conditions. (**F**) Quantification of the ratio of globular to filamentous actin fluorescent intensities. (**G**) Scanning electron microscopy of filament network between groups. Top panel is the lowest magnification with higher magnifications in panels (i) and (ii). Bottom-cartoon representation of SEM filament network and hypothesized role of Rab35. * p < 0.05, ** p<0.01, **** p < 0.0001, NS=Non-Significant. Error bars represent 95% confidence intervals. Insets are areas of higher magnification. All experiments were done using Human umbilical vein endothelial cells in triplicate.

Based off this finding, we reasoned that if Rab35 was involved with actin polymerization, then knockdown of Rab35 would shift the balance between globular and filamentous actin to skew more globular, as less filaments are being assembled. Using differential centrifugation, we separated the globular and filamentous pools of actin as previously reported [52]. Rab35 knockdown significantly increased the globular actin abundance compared with control (**Fig. 6C,D**). Using a similar method, we stained for globular actin using GC globulin and phalloidin to detect the filamentous actin [53]. Again, our results demonstrated an increase in globular to filamentous actin ratio in the absence of Rab35 as compared with controls (**Fig. 6E,F**). We also co-stained for globular and filamentous actin while expressing Rab35 to ensure Rab35 colocalized with both actin populations. Indeed, Rab35 was strongly localized to sites of globular actin that were also positive for filamentous actin (**Fig. S8F**).

Lastly, we used scanning electron microscopy to better visualize the remaining actin network in ECs depleted of Rab35 or treated with CK-666. Qualitatively, there was reduced filament density in the lamellipodia regions of the Rab35 depleted and CK-666 treated conditions as compared with control (**Fig. 6G**). In Rab35 depleted ECs, we also observed elevated instances of bundles of actin that were more disorganized in appearance as compared with control (**Fig. 6G**), potentially representing a compensatory effect for the lack of filamentous actin. Overall, these results suggest that Rab35 is associated with regulating local sites of actin assembly.

### Rab35 is required for blood vessel development in zebrafish

We next generated a Rab35 knockout in zebrafish using CRISPR/Cas9 gene editing to test if Rab35 was also required for in vivo angiogenic processes [54]. In zebrafish, we targeted both Rab35 paralogs, Rab35A and Rab35B. By sequence analysis we observed 100% indel formation in F_0_ injected zebrafish for both Rab35 paralogs (**Fig. 7A**). Double Rab35A/B knockout was embryonic lethal marked by a lack of normal development as compared with scramble guide injected controls, suggesting Rab35 is critical for normal embryonic development (**Fig. 7B**). However, we did see a spectrum of developmental defects when the single-guide RNA amount were diluted. In a vascular Lifeact-GFP expressing line injected with a sublethal dosage of Rab35A/B single-guide RNA, we focused on actin defects. Here, we did not quantify vascular defects due to the generalized tissue dysmorphogenesis of these embryos; alternatively, our goal was to determine if similar actin accumulations occurred in vivo as observed in vitro. In line with our in vitro data, we observed a significant increase in actin aggregations in the Rab35A/B knockout group compared with controls (**Fig. 7C,D**). Similarly, overexpression of the DN Rab35 mutant or treatment with CK-666 promoted an increase in aberrant Rab35 accumulations, presumably bound to actin (**Fig. 7E,F**). To subvert the lethality of global Rab35A/B deletion, we generated chimeric embryos using blastomere transplants [55]. Transfer of Rab35A/B CRISPR injected cells into a WT host generated mosaic intersomitic blood vessels (ISVs) allowing for comparison of both WT and Rab35A/B null blood vessels side-by-side. Similar to in vitro results, Rab35A/B null ISVs were dysmorphic, marked by a thin appearance and the absence of a lumen as assessed by microangiography (**Fig. 7G,H**). Overall, these results indicate Rab35 is necessary for organismal viability and actin homeostasis in vivo.

**Figure 7.**
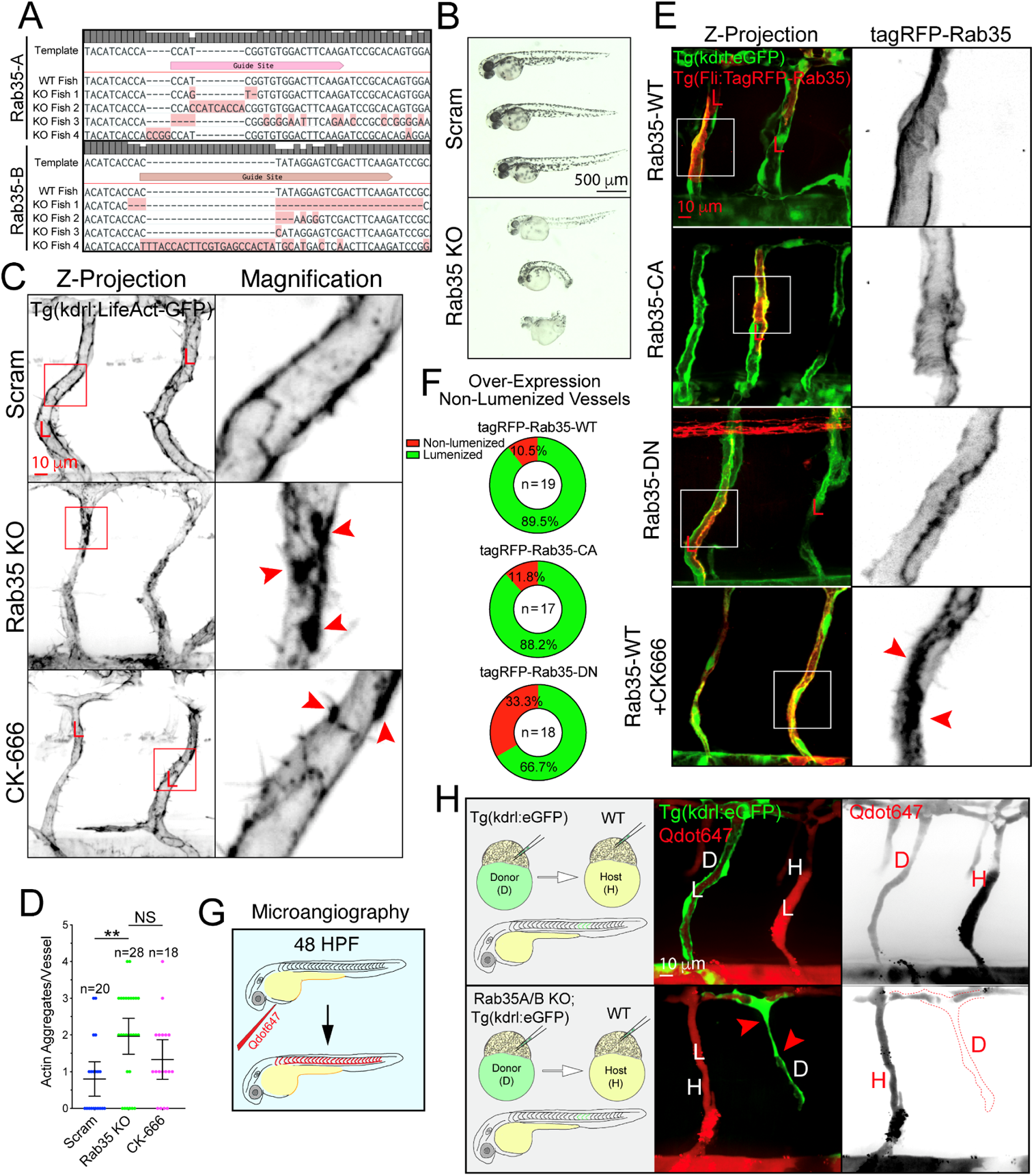
Rab35 is required for blood vessel development in zebrafish. (**A**) CRISPR-mediated knockout of Rab35A/B confirmation by sequencing. Four random fish were sequenced following CRISPR/guide injections. (**B**) Zebrafish morphology at 48 hours post fertilization (hpf) post injection of scramble (Scram) and Rab35A/B CRISPR guides. (**C**) Representative images of intersomitic blood vessels (ISVs) of Scram and Rab35A/B knockout as well as CK-666 (Arp Inhibitor) treated zebrafish at 48 hpf expressing endothelial specific LifeAct-GFP. Red arrowheads indicate abnormal aggregates of actin. (**D**) Quantification of actin aggregates between groups. N= number of ISVs. A minimum of 5 fish were used per group. (**E**) Representative images of mosaic expression of Tag-RFP-Rab35 WT (top row), CA (second row), DN (third row) and WT with CK-666 treatment in zebrafish at 48 hpf. Red arrowheads depict excess of Rab35 at the plasma membrane. (**F**) Quantification of non-lumenized vessels at 48 hpf between groups mentioned in panel E. (**G**) Cartoon representation of microangiography in zebrafish larvae using quantum dots 647 (Qdot647) at 48 hpf. (**H**) Representative images of ISVs after transplantation of Tg(kdrl:GFP) donor (D) into Tg(kdrl:mCherry) host (H) (top panels). Bottom panels-representative images of ISVs after transplantation of Rab35A/B knockout donor cells from Tg(kdrl:GFP) line into Tg(kdrl:mCherry) host. Red arrowheads indicate lumen failure. ** p<0.01, NS=Non-Significant. Error bars represent 95% confidence intervals. Insets are areas of higher magnification. All experiments were done in triplicate.

## DISCUSSION

In the current work, we explored the contribution of Rab35 to angiogenic sprouting behaviors vital to blood vessel development. The primary goal of this work was to interpret what of the many reported functions of Rab35 matters most during blood vessel morphogenesis by systematically characterizing Rab35 itself and the downstream effector pathways. Using a combination of 3D sprouting, biochemistry and in vivo gene editing, we demonstrate that Rab35’s most prominent function is to regulate actin dynamics during angiogenesis. More specifically, we show that the GEF DENNd1c tethers active Rab35 to the actin cytoskeleton. Once localized to actin, Rab35 promotes actin polymerization and remodeling required for sprout formation. Additionally, we confirmed that Rab35 is required for blood vessel development in zebrafish. To our knowledge, this is the first investigation demonstrating the requirement of Rab35 for blood vessel function and the first investigation in any tissue dissecting Rab35’s most dominant biological role accounting for the most prominent effector pathways.

The genesis of the current project was originally aimed to characterize how podocalyxin was trafficking in ECs, as this is still an outstanding question in the field. Our past work demonstrated Rab27a, that was largely implicated in podocalyxin trafficking in epithelial cells, was not related to this pathway [7], thus our very next candidate was Rab35. Others have comprehensively established a direct association between Rab35 and podocalyxin as well as the downstream impact on lumen biogenesis [15, 16]. Our data in the current investigation once again shows that endothelial trafficking signatures greatly differ from epithelial programs. More specifically, we expansively tested for both localization and direct binding interaction between Rab35 and podocalyxin of which we found none. However, this negative result prompted us to further investigate Rab35 function during angiogenic sprouting.

Rab proteins are the most numerous subset of Ras family small guanosine triphosphatases (GTPases). Rab proteins control biogenesis, movement, and docking of vesicles in specific trafficking pathways by recruiting unique effector proteins to different membrane compartments [13, 56]. Rab35, in particular, has been shown to have many roles that vary by tissue type, organism, and developmental stage. In distilling the literature, it can be argued that Rab35 has four major effectors that mediate its function in vertebrates: RUSC, MICAL-L1, ACAP2, and OCRL. Given Rab-family GTPases exert their function via effector interaction, we began by first establishing that Rab35 was required for sprouting, and then determined how each effector contributed to the loss of Rab35 phenotype. Surprisingly, RUSC, MICAL-L1 and OCRL either showed no phenotypic contribution to sprouting or failed to directly bind Rab35. The most promising candidate ACAP2 exhibited the best phenotype for recapitulating the Rab35 loss of function effect. In terms of ACAP2, we had several interesting findings. The predominant hypothesis is that GTP Rab35 binds ACAP2 sequestering its ability to inactivate Arf6, resulting in a gain of function for Arf6. In ECs, we could not confirm direct binding between ACAP2 and Arf6, we also did not observe that Rab35 knockdown affected Arf6 activity as previously reported [17–19, 29, 32, 33, 37, 38]. These previous reports were carried out in non-endothelial tissue, which may explain the signaling discrepancy. However, we also overexpressed a CA Arf6, the predicted outcome of loss of Rab35 in the aforementioned epithelial systems, and also could not phenocopy the Rab35 loss of function effect on sprouting, again suggesting this signaling pathway is not essential in ECs in the absence of Rab35.

A major finding was that the GEF DENNd1c played a key role in Rab35 function. Canonically, GEFs primarily convert proteins from a GDP to GTP-bound state; however, DENNd1c is evolutionarily divergent from both DENNd1a/b that solely control Rab35 GTPase activity [41]. In our hands, loss of DENNd1c did not alter GTP activation, but controlled the localization of Rab35 to actin fibrils. Knockdown of DENNd1c strongly phenocopied loss of Rab35 suggesting that localization to actin is a primary function of Rab35 during sprouting angiogenesis.

Actin plays a pivotal role in angiogenesis both from a cell migration and vessel stabilization aspect [57–62]. Loss of normal actin architecture has been shown to drastically affect virtually all facets of blood vessel formation [63–67]. In this sense, our results are not surprising in that actin misregulation promoted such a profound effect on sprouting parameters. However, given Rab35’s broad scope of function as well as never being characterized in angiogenic processes, it would be exceedingly hard to predict. Moreover, actin regulation is typically known to be directly controlled through more conventional signaling paradigms such as Rac1 and CDC42. Our results paint a novel scenario that trafficking-based regulators can control vital crosstalk with the actin cytoskeleton. It is still an outstanding question what of the hundreds of cytoskeletal proteins Rab35 is interfacing with to participate in actin regulation processes.

Overall, our investigation is the first to systematically rule out other known Rab35 pathways, highlighting Rab35’s novel function in mediating actin dynamics during blood vessel formation in vitro and in vivo. We believe this work is important not only from the vantage of understanding EC biology and its unique trafficking signatures, but from a disease standpoint as Rab35 is commonly upregulated in solid cancers [68]. In general, we contend that mapping endothelial trafficking patterns will shed important light on how ECs orchestrate blood vessel formation by integrating both cell-autonomous and collective-cell signaling.

## ACKNOWLEDGEMENTS

Work was supported by funding from the National Heart Lung Blood Institute (Grant 1R56HL148450-01, R15HL156106-01A1, R01HL155921-01A1) (EJK).

## CONTRIBUTIONS

CRF, HK and EJK performed all experiments. CRF and EJK wrote the manuscript.

## SUPPLEMENTAL FIGURES

**Supplemental Figure 1.**
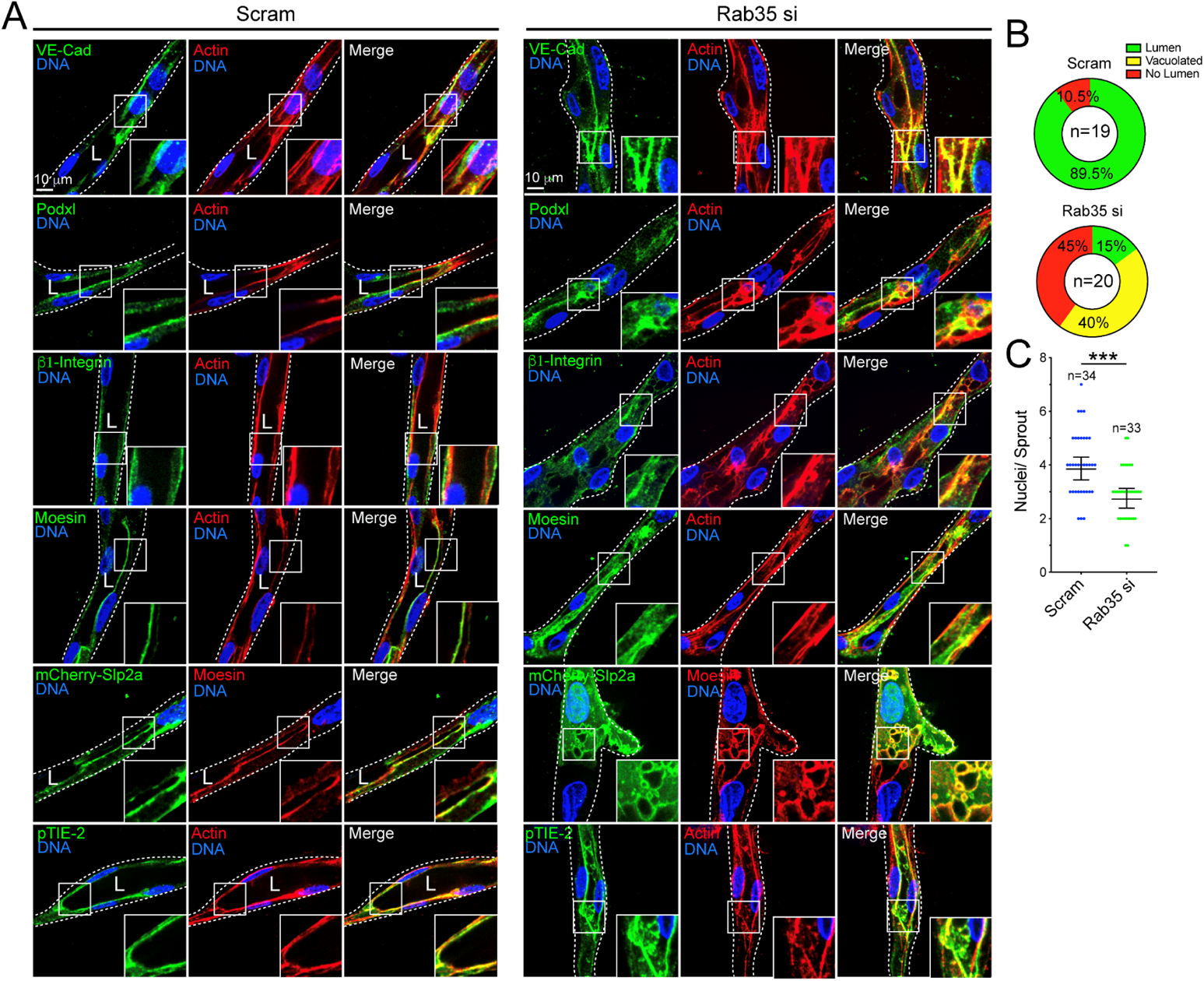
Knockdown of Rab35 distorts cell apicobasal polarity. (**A**) Scramble (Scram) and Rab35 siRNA(si)-treated sprouts stained for VE-cadherin (VE-cad), podocalyxin (Podxl), β1-integrin, moesin or phosphorylated Tie2 (pTie2) apical and basal protein markers. Apical marker synaptotagmin-like protein 2a (mCherry-Slp2a) was transduced into sprouts. L denotes lumen and white dotted lines outline sprout exterior. (**B**) Quantification of lumen formation in Scram and Rab35 siRNA-treated sprouts. Lumens were defined as an open continuous cavity. Vacuolated sprouts were defined as sprouts lacking a contiguous lumen, while exhibiting an excess of large vacuoles. The no lumen group was defined as sprouts that had no visible cavity or vacuoles. N=number of sprouts. (**C**) Quantification of nuclei per sprout in Scram and Rab35 siRNA treated sprouts. *** p<0.001. Error bars represent 95% confidence intervals. Insets are areas of higher magnification. All experiments were done using Human umbilical vein endothelial cells in triplicate.

**Supplemental Figure 2.**
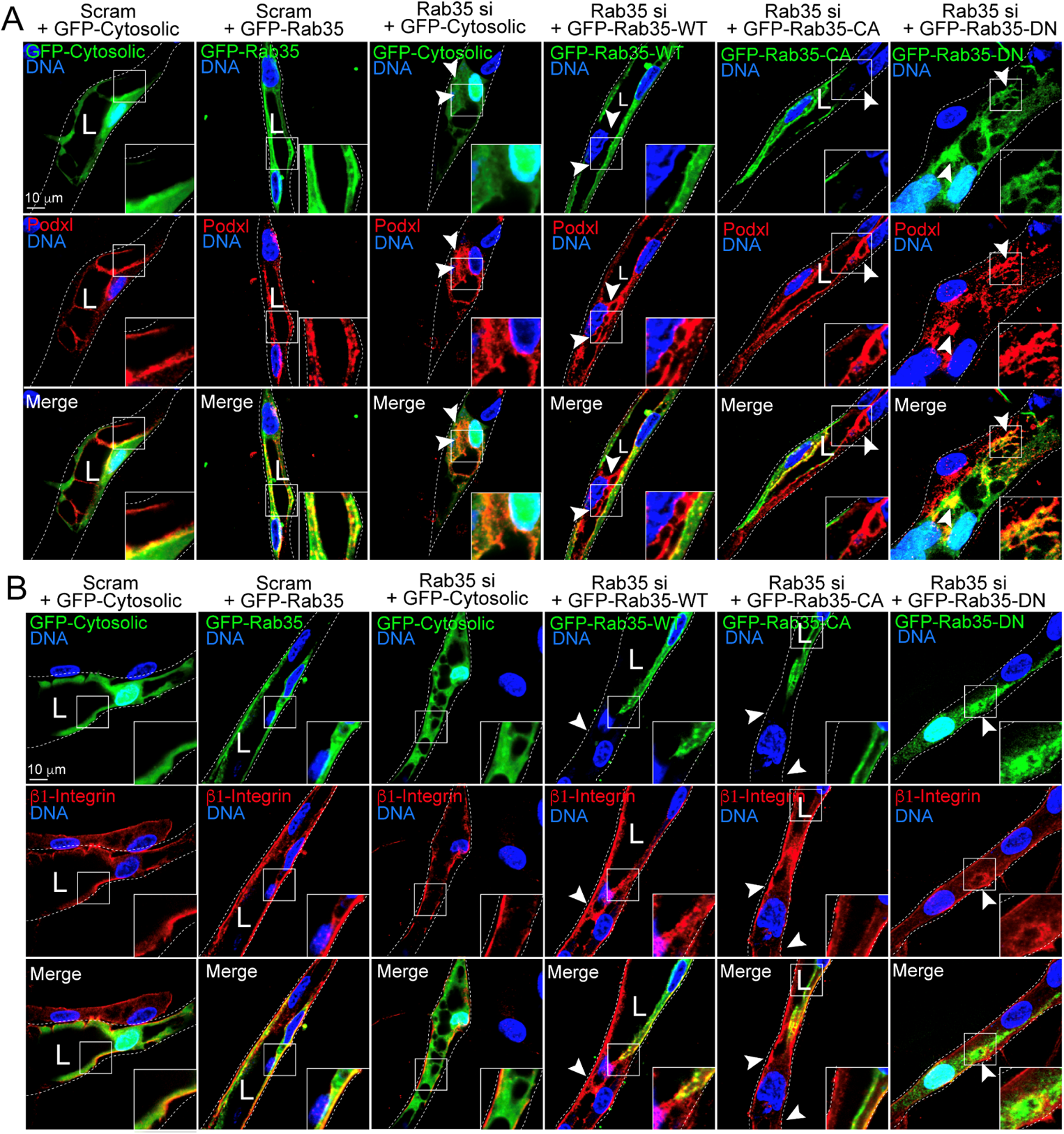
Rab35 knockdown disrupts sprout polarity programs. (**A,B**) Representative images of Rab35 knockdown (KD) sprouts transfected with cytosolic GFP or GFP-Rab35 wild type (WT), constitutively-active (CA) or dominant negative (DN) for rescues. Sprouts were also stained for apical marker podocalyxin (Podxl) or apical marker β1-integrin. Arrowheads denote abnormal localization of podocalyxin or β1-integrin. L denotes lumen in all images. White dotted lines mark sprout exterior. Insets are areas of higher magnification. All experiments were done using Human umbilical vein endothelial cells in triplicate.

**Supplemental Figure 3.**
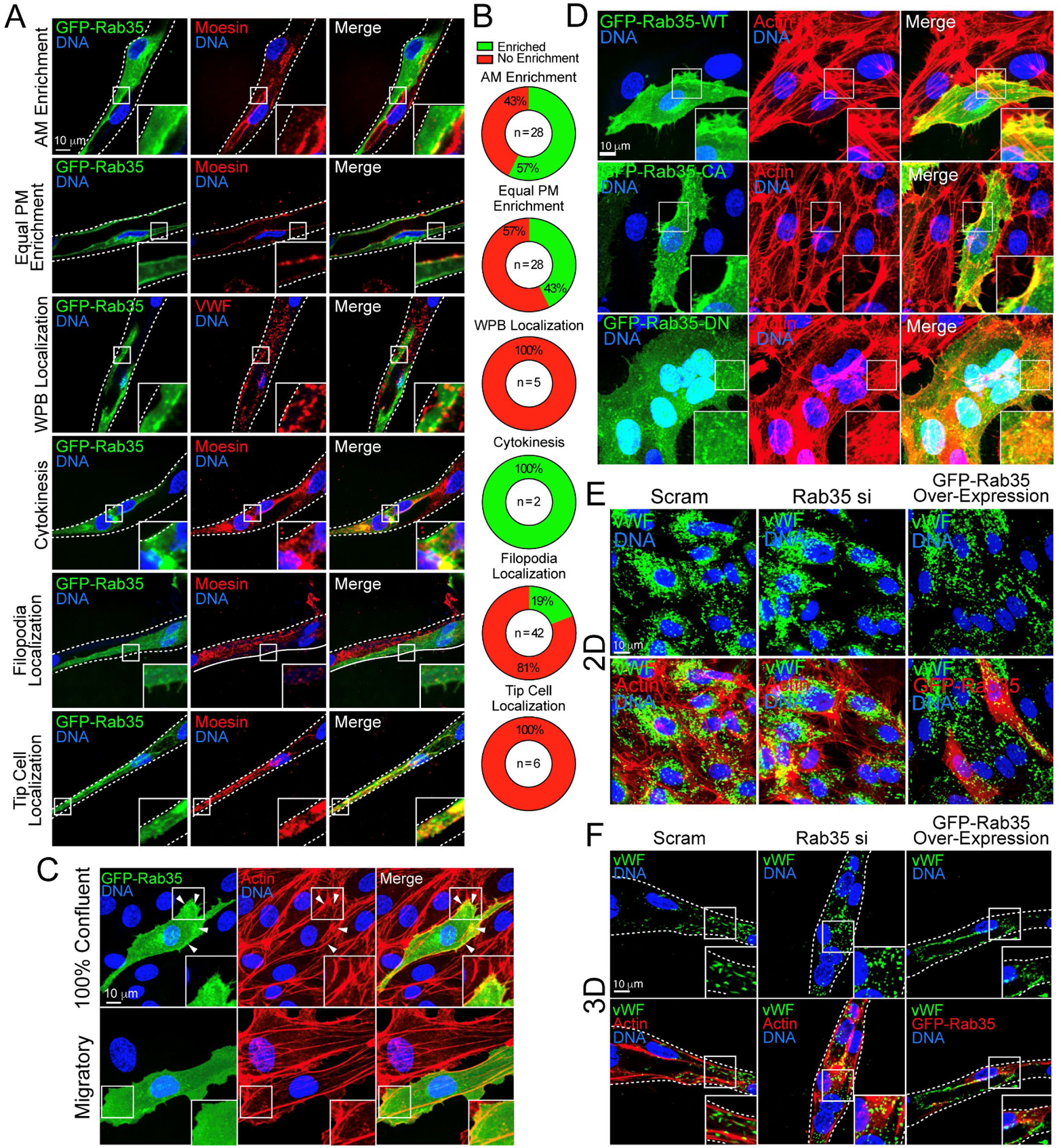
Rab35 localizes to the plasma membrane and not to Weibel-Palade Bodies. (**A**) Representative images of GFP-Rab35 localization binned by its proximity to the apical plasma membrane (AM), equal enrichment at the basal and apical plasma membrane (equal plasma membrane (PM) enrichment), Weibel-Palade bodies (WPBs), at sites of cytokinesis, filopodia, and most distal cell in the sprout (tip cell). Sprouts were also stained for moesin to mark the apical membrane. (**B**) Quantification of GFP-Rab35 enrichment with respect to the described conditions in panel A. (**C**) Representative images of GFP-Rab35 localization in 2-dimensional culture stain for actin. The top panels are of a confluent monolayer and the bottom panels are of migratory sub-confluent cells. Arrowheads indicate co-localization of actin and GFP-Rab35. (**D**) Representative images of 2-dimensional localization of GFP-Rab35 wild type (WT, top panels), constitutively active (CA, middle panels), and dominant negative (DN, bottom panels) stained for actin. (**E**) Representative images of cells treated with scramble (Scram) or Rab35 siRNA (si) and stained for WPB marker von Willebrand Factor (vWF) and actin or overexpressing GFP-Rab35. (**F**) Representative images of sprouts treated with Scram or Rab35 siRNA stained for vWF and actin or expressing GFP-Rab35 in 3-dimensional (3D) sprouts. Insets are areas of higher magnification. White dotted lines mark sprout exterior. All experiments were done using Human umbilical vein endothelial cells in triplicate.

**Supplemental Figure 4.**
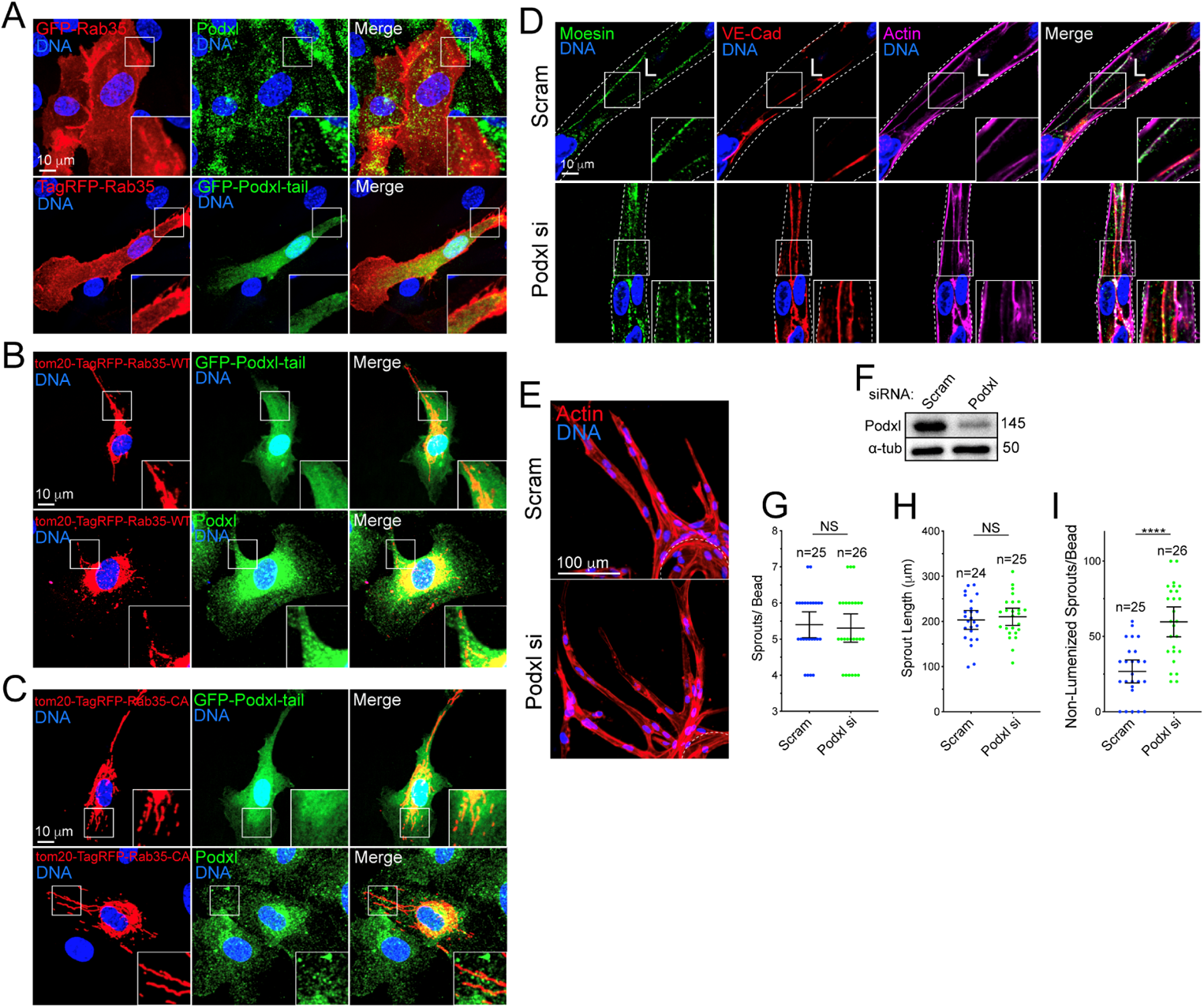
Rab35 does not affect podocalyxin trafficking. (**A**) Two-dimensional localization of GFP-Rab35 with podocalyxin (Podxl) (top panels) and GFP-Podxl-tail (bottom panels). (**B**). Top panels-cell co-expressing tom20-TagRFP-Rab35-wild type (WT) with GFP-Podxl-tail. Bottom panels-cell expressing tom20-TagRFP-Rab35-WT stained for endogenous podocalyxin. (**C**) Representative image of a cell co-expressing tom20-TagRFP-Rab35-constitutively active (CA) mutant with GFP-Podxl-tail. Bottom panels show a cell expressing tom20-tagRFP-Rab35-constitutively active (CA) mutant stained for endogenous podocalyxin. (**D**) Representative image of sprouts treated with scramble (Scram) or podocalyxin siRNA (si) and stained for moesin, VE-cadherin (VE-cad) and actin. L denotes lumen. White dotted lines mark sprout exterior. (**E**) Sprout morphology for the same conditions as D. (**F**) Confirmation of siRNA-mediated knockdown by western blot. (**G-I**) Quantification of indicated sprouting parameters across groups. **** p < 0.0001, NS=Non-Significant. Error bars represent 95% confidence intervals. Insets are areas of higher magnification. All experiments were done using Human umbilical vein endothelial cells in triplicate.

**Supplemental Figure 5.**
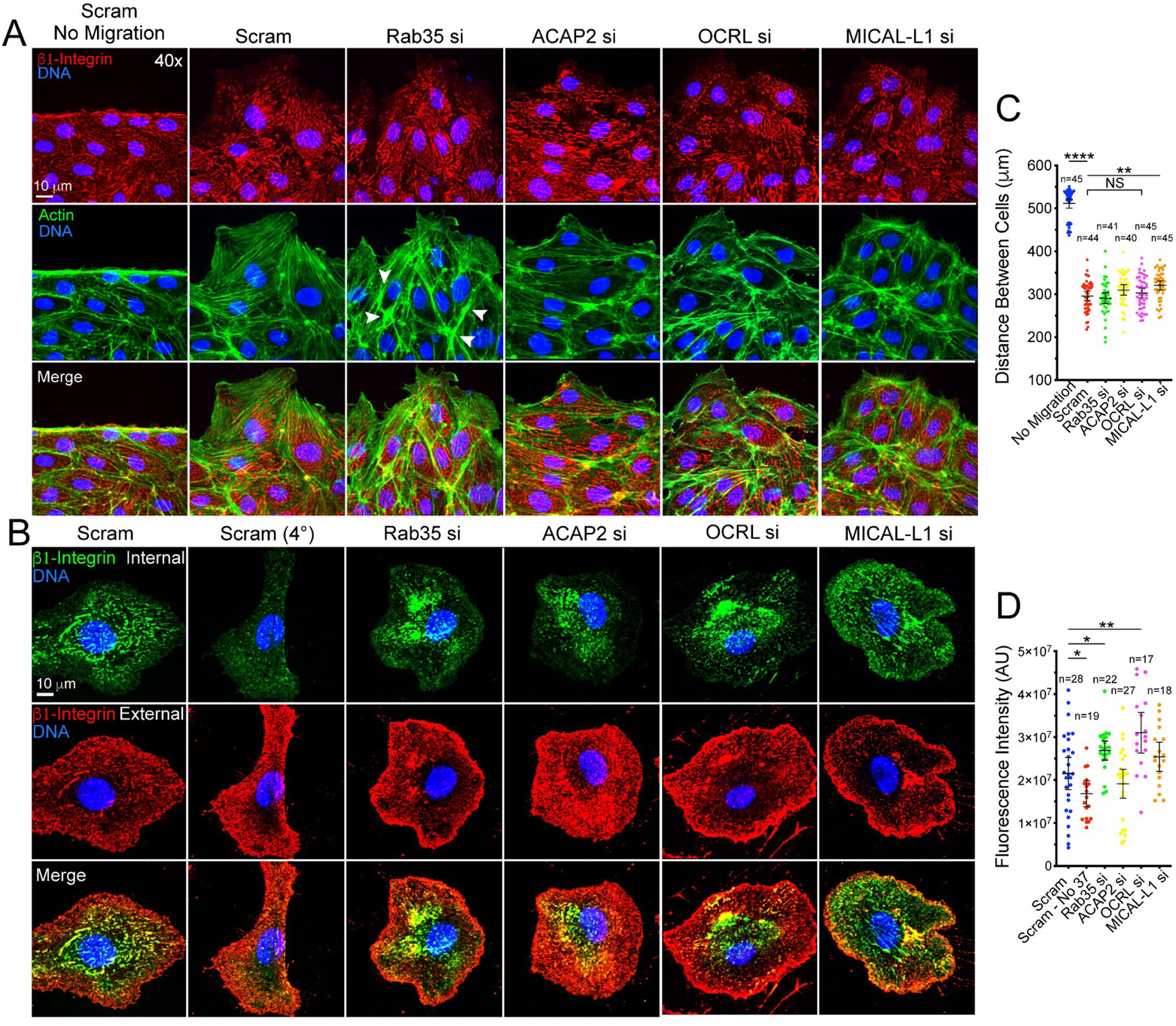
Knockdown of Rab35 impacts integrin internalization, but not cell migration. (**A**) Migration assay in cells treated with scramble (Scram), Rab35, ACAP2, OCRL, or MICAL-L1 siRNA (si). Cells were stained for β1-integrin and actin. Arrowheads indicate abnormal actin architecture, namely elevated actin deposition. (**B**) Antibody feeding assay to test for integrin turnover between conditions. Cells were treated with indicated siRNA. Green channel represents internalized integrins, while the red channel marks only external integrins. As a control to inhibit endocytosis a group was held at 4°C. (**C**) Quantification for the migration assay in A. (**D**) Fluorescence intensity of internalized β1-integrin in panel B. * p<0.05, ** p<0.01, **** p < 0.0001, NS=Non-Significant. Error bars represent 95% confidence intervals. All experiments were done using Human umbilical vein endothelial cells in triplicate.

**Supplemental Figure 6.**
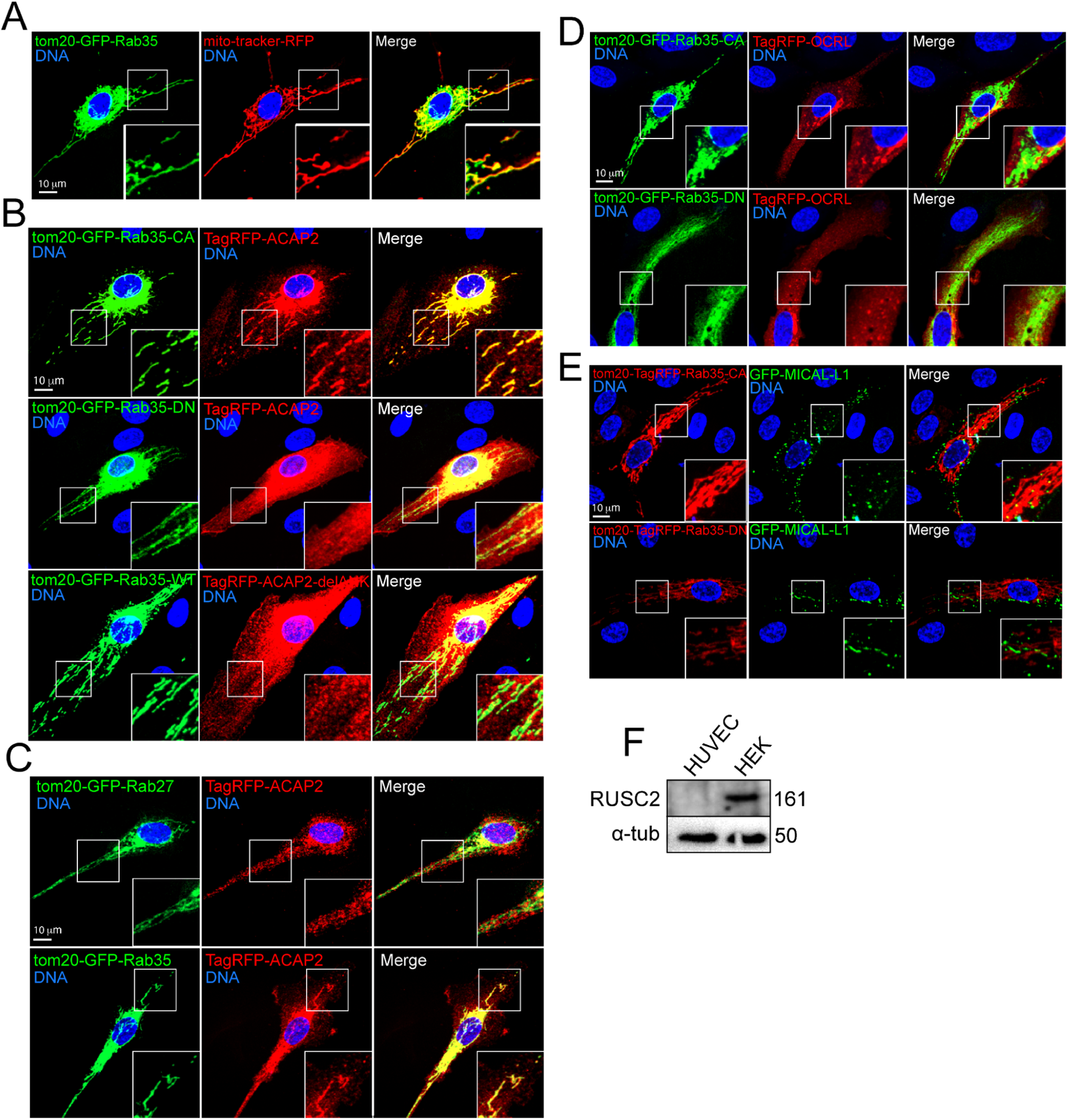
Rab35 binds only ACAP2. (**A**) Cells stained for mitochondria (Mito-tracker) and transfected with tom20-tagRFP-Rab35. (**B**) Representative images of a cell co-expressing tom20-tagRFP-Rab35-wild type (WT), constitutively active (CA), or dominant negative (DN) variants with TagRFP-ACAP2 or ACAP2 with deleted ankyrin repeat domain (delANK). (**C**) Representative image of a cell expressing tagRFP-ACAP2 and tom20-GFP-Rab27-WT (top panels). Bottom panel is a representative image of a cell expressing of tom20-GFP-Rab35-WT with tagRFP-ACAP2. (**D**) Top panels-representative image of a cell expressing tom20-GFP-Rab35-CA and OCRL. Bottom panels-cell expressing tom20-GFP-Rab35-DN and TagRFP-OCRL. (**E**) Top panels-representative image of a cell expressing tom20-TagRFP-Rab35-CA and GFP-MICAL-L1. Bottom panels-cell expressing tom20-TagRFP-Rab35-DN and GFP-MICAL-L1. (**F**) Western blot image probing for RUSC2 in both HEK293 cells and Human umbilical vein endothelial cells (HUVECs). Insets are areas of higher magnification. All experiments were done using Human umbilical vein endothelial cells in triplicate.

**Supplemental Figure 7.**
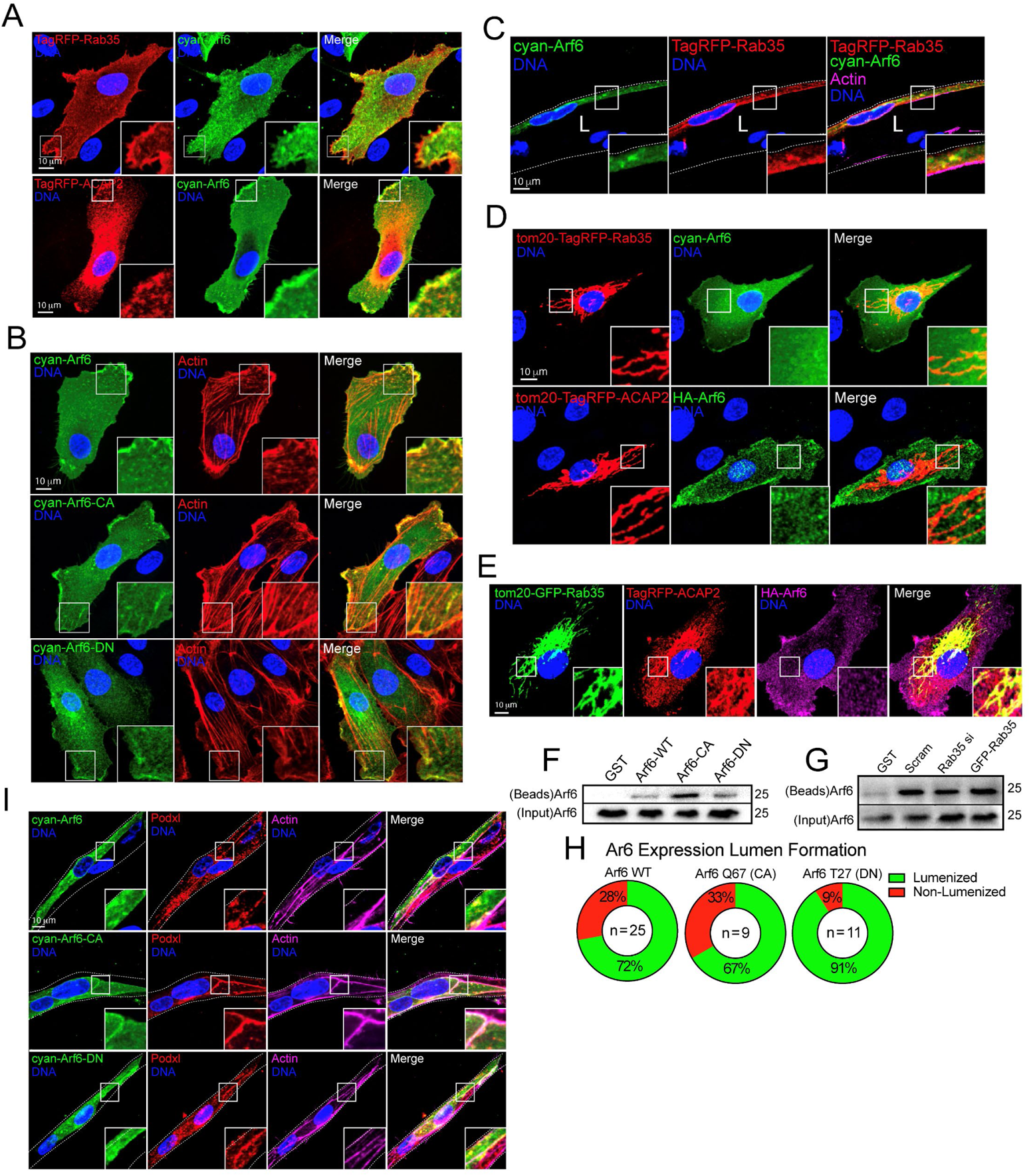
Rab35 does not affect Arf6 activity in endothelial cells. (**A**) Two-dimensional localization of cyan-Arf6 with tagRFP-Rab35 (top panels) and ACAP2 (bottom panels). (**B**) Two-dimensional localization of cyan-Arf6-wild type (WT, top panels), constitutively active (CA, middle panels), and dominant negative (DN, bottom panels) stained for actin. (**C**) Localization of tag-RFP-Rab35 and cyan-Arf6 in a sprout. (**D**) Top panel-representative image of a cell expressing tom20-tagRFP-Rab35 and cyan-Arf6. Bottom panel-representative image of a cell expressing tom20-tagRFP-ACAP2 and HA-Arf6. (**E**) Representative image of a cell expressing tom20-GFP-Rab35, tagRFP-ACAP2 and HA-Arf6. (**F**) Pulldown assay using GGA3 to probe for activated Arf6. Cells were transfected with WT, CA, or DN Arf6. (**G**) Pulldown assay using GGA3 to probe for activated Arf6. Cells were treated with scramble (Scram) and Rab35 siRNA (si) or transfected with GFP-Rab35. () Quantification of open or collapsed lumens after transfection with WT, CA, or DN cyan-Arf6. N= number of sprouts. (**I**) Representative images of sprouts transduced with WT, CA, or DN cyan-Arf6 stained for Podocalyxin (Podxl) and actin. L denotes lumen in all images. White dotted lines mark sprout exterior. Insets are areas of higher magnification. All experiments were done using Human umbilical vein endothelial cells in triplicate.

**Supplemental Figure 8.**
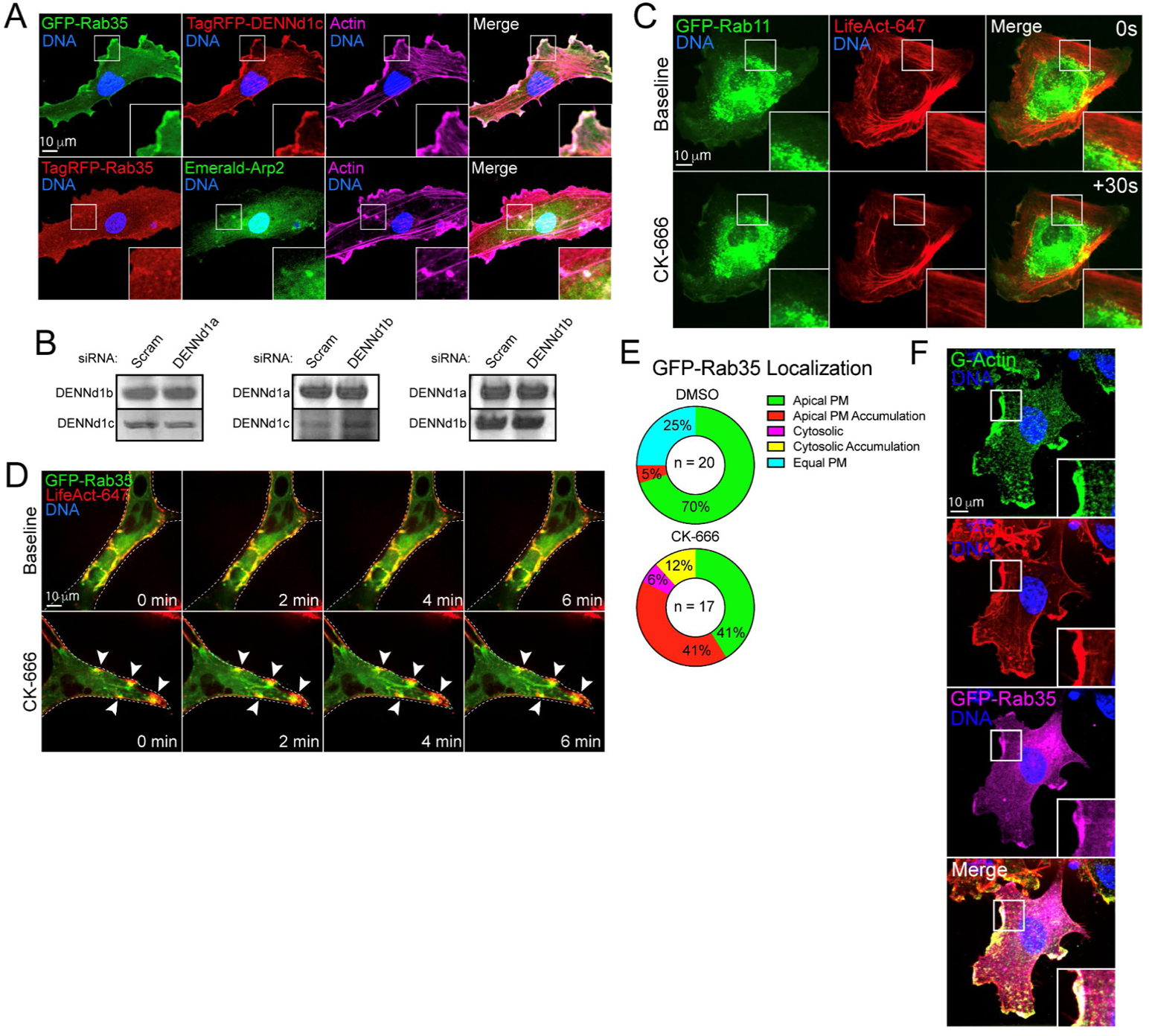
Rab35 is recruited to sites of active actin polymerization. (**A**) Top panel-representative image of a cell expressing GFP-Rab35 and tagRFP-DENNd1c. Bottom panel-representative image of a cell expressing GFP-Rab35 and Emerald-Arp2. (**B**) Western blot of DENNd1a-c knockdown. For each blot a DENNd1 was knocked down and the remaining two DENND1s were probed for to test for compensation effects. (**C**) Representative live-image of a cell expressing GFP-Rab11 and TagRFP647 (647)-LifeAct before and after CK-666 treatment. (**D**) GFP-Rab35 and LifeAct-647 co-expression in sprout live-imaged at baseline and following treatment with CK-666. Arrowheads indicate accumulations of GFP-Rab35 and LifeAct-647. Dotted line indicates sprout exterior. (**E**) Quantification of GFP-Rab35 localization upon DMSO (vehicle) or CK-666 administration. Apical plasma membrane (PM, uniformly localized to apical membrane), apical PM accumulation (Rab35 puncta at the apical membrane), cytosolic (localized in the cytoplasm), cytosolic accumulations (Rab35 puncta in the cytoplasm), equal PM (Rab35 equally distributed between the apical and basal membranes). Two-dimensional localization of GFP-Rab35 with globular-actin and filamentous-actin. (**F**) Representative image of a cell expressing GFP-Rab35 and stained for filamentous (F) and globular (G) actin. Insets are areas of higher magnification. All experiments were done using Human umbilical vein endothelial cells in triplicate.

## MOVIE FIGURE LEGENDS

**Movie 1.** Sprout expressing GFP-Rab35. L denotes lumen.

**Movie 2.** Sprout expressing GFP-Rab35 and LifeAct-TagRFP647 (FarRed) treated with scrambled siRNA. L denotes lumen.

**Movie 3.** Sprout expressing GFP-Rab35 and LifeAct-TagRFP647 (FarRed) treated with DENNd1c siRNA. L denotes lumen. Arrow marks actin accumulation.

**Movie 4.** Sprout expressing GFP-Rab35 and LifeAct-TagRFP647 (FarRed) treated with DMSO. Arrow marks normal actin buttressing at junctions.

**Movie 5.** Sprout expressing GFP-Rab35 and LifeAct-TagRFP647 (FarRed) treated with CK-666. Arrow marks actin accumulations.

**Movie 6.** Cell expressing TagRFP-Rab35 and GFP-LifeAct before and after CK-666 treatment.

**Movie 7.** Cell expressing TagRFP-DENNd1c and GFP-LifeAct before and after CK-666 treatment.

**Movie 8.** Cell expressing mCherry-Arp2 and GFP-Rab35 before and after CK-666 treatment.

**Movie 9.** Cell expressing GFP-Rab35, LifeAct-TagRFP647 (FarRed), and ligand-modulated antibody fragments targeted to the mitochondria (mito-LAMA) before and after trimethoprim (TMP) administration.

**Movie 10.** Cell expressing GFP-Rab35, mCherry-Arp2, and ligand-modulated antibody fragments targeted to the mitochondria (mito-LAMA) before and after trimethoprim (TMP) administration.

**Movie 11.** Cell expressing GFP-Rab35, TagRFP-DENNd1c, and ligand-modulated antibody fragments targeted to the mitochondria (mito-LAMA) before and after trimethoprim (TMP) administration.

**Movie 12.** Cell expressing GFP-Rab35, mCherry-Arp2, and ligand-modulated antibody fragments targeted to the mitochondria (mito-LAMA) before and after trimethoprim (TMP) administration. After TMP treatment cell were also treated with CK-666.

**Movie 13.** DENNd1c knockdown (siRNA) cell expressing GFP-Rab35, mCherry-Arp2, and ligand-modulated antibody fragments targeted to the mitochondria (mito-LAMA) before and after trimethoprim (TMP) administration.

## MAJOR RESOURCE TABLE

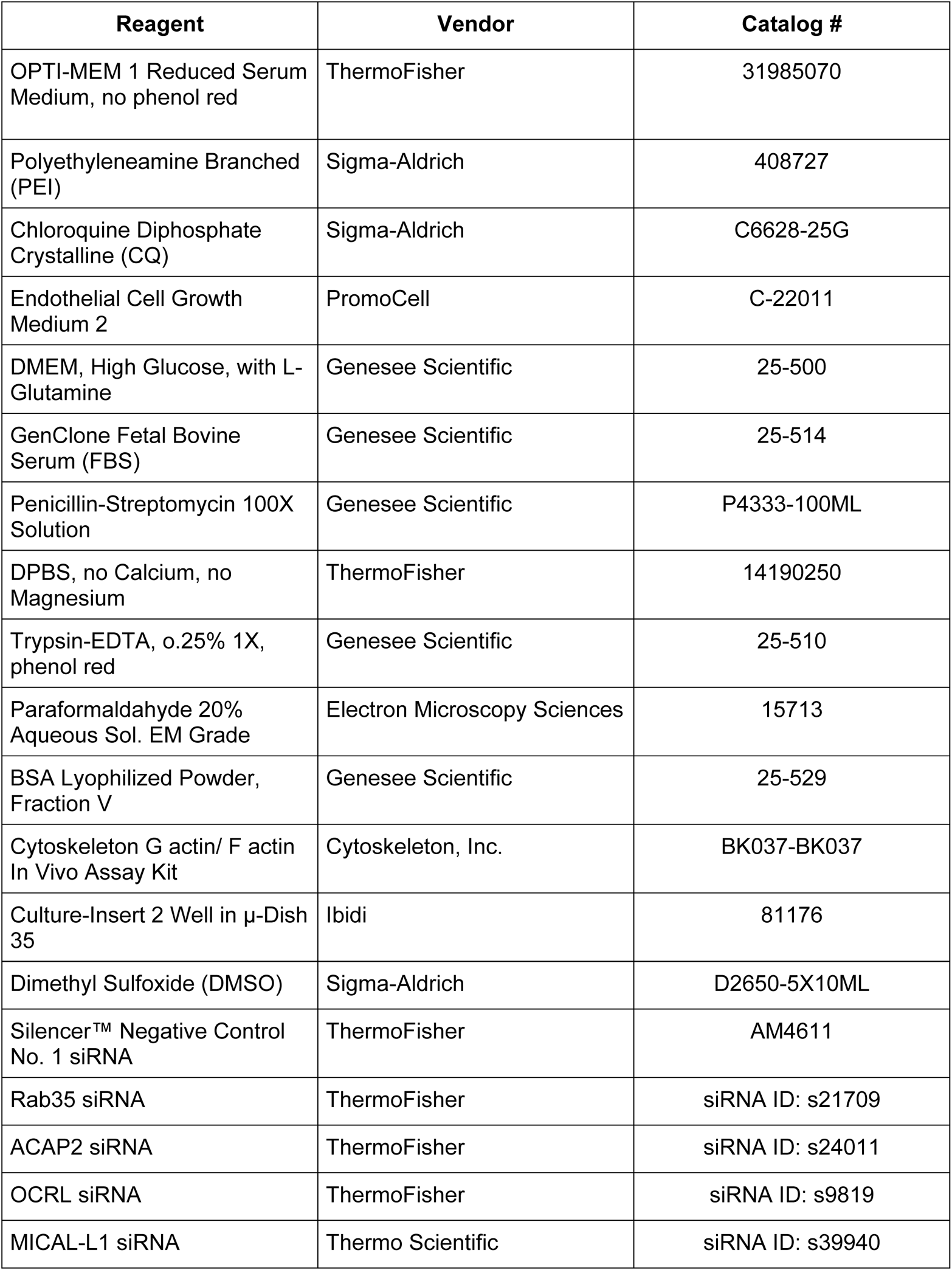
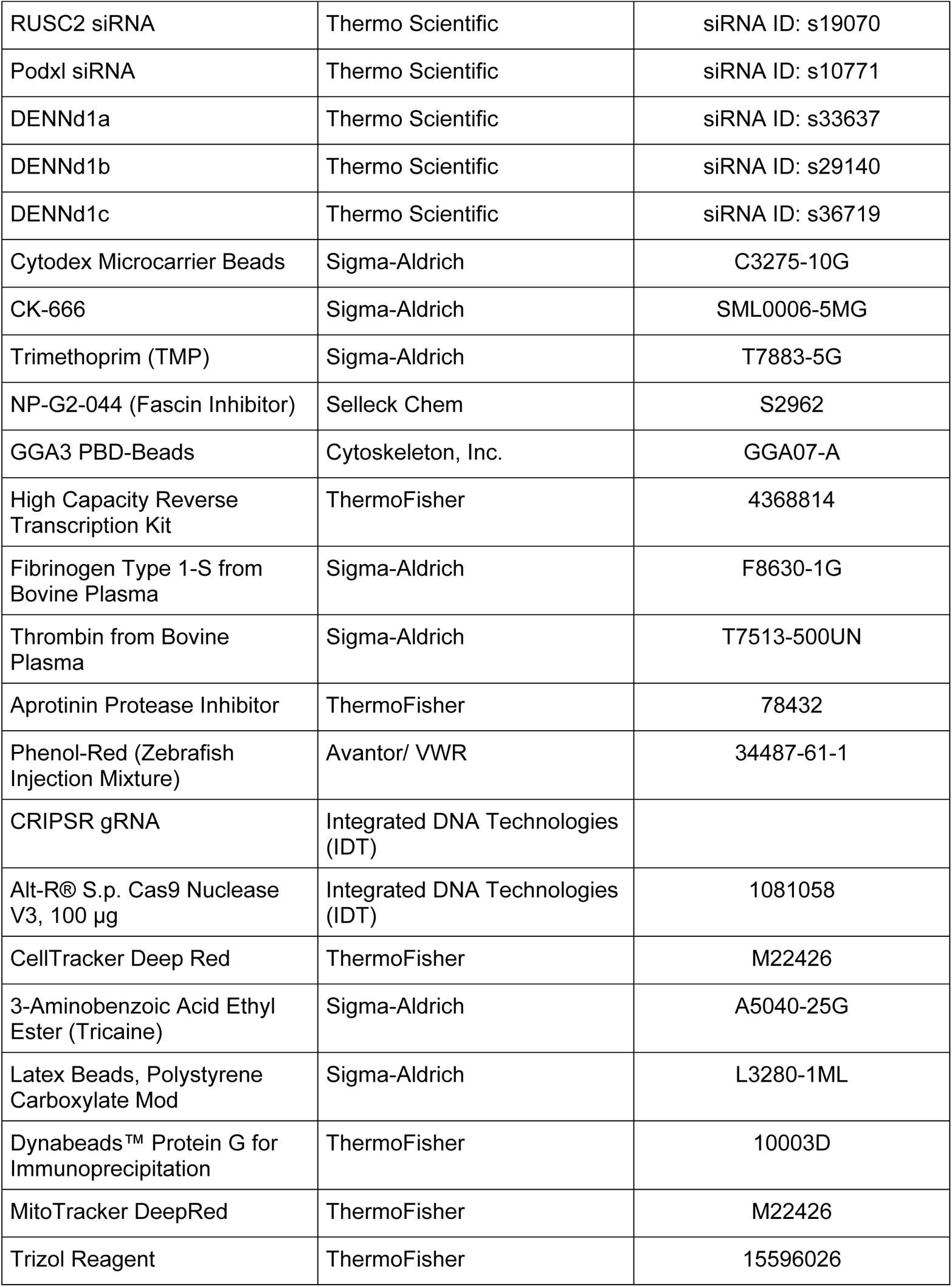
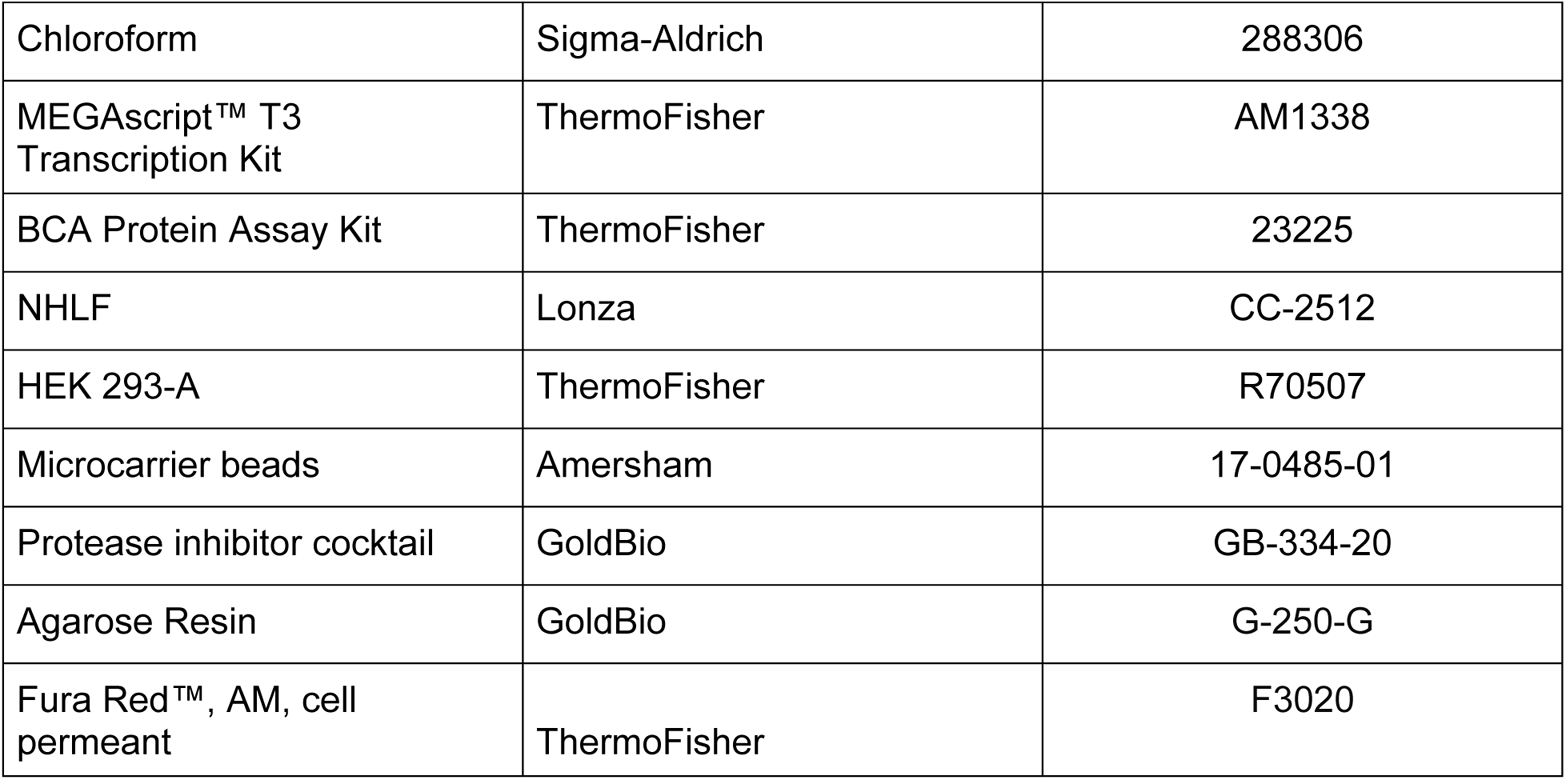

| **Reagent** | **Vendor** | **Catalog #** |
| --- | --- | --- |
| OPTI-MEM 1 Reduced Serum Medium, no phenol red | ThermoFisher | 31985070 |
| Polyethylenamine Branched (PEI) | Sigma-Aldrich | 408727 |
| Chloroquine Diphosphate Crystalline (CQ) | Sigma-Aldrich | C6628-25G |
| Endothelial Cell Growth Medium 2 | PromoCell | C-22011 |
| DMEM, High Glucose, with L- Glutamine | Genesee Scientific | 25-500 |
| GenClone Fetal Bovine Serum (FBS) | Genesee Scientific | 25-514 |
| Penicillin-Streptomycin 100X Solution | Genesee Scientific | P4333-100ML |
| DPBS, no Calcium, no Magnesium | ThermoFisher | 14190250 |
| Trypsin-EDTA, 0.25% 1X, phenol red | Genesee Scientific | 25-510 |
| Paraformaldehyde 20% Aqueous Sol. EM Grade | Electron Microscopy Sciences | 15713 |
| BSA Lyophilized Powder, Fraction V | Genesee Scientific | 25-529 |
| Cytoskeleton G actin/ F actin In Vivo Assay Kit | Cytoskeleton, Inc. | BK037-BK037 |
| Culture-Insert 2 Well in μ-Dish 35 | Ibidi | 81176 |
| Dimethyl Sulfoxide (DMSO) | Sigma-Aldrich | D2650-5X10ML |
| Silencer™ Negative Control No. 1 siRNA | ThermoFisher | AM4611 |
| Rab35 siRNA | ThermoFisher | siRNA ID: s21709 |
| ACAP2 siRNA | ThermoFisher | siRNA ID: s24011 |
| OCRL siRNA | ThermoFisher | siRNA ID: s9819 |
| MICAL-L1 siRNA | Thermo Scientific | siRNA ID: s39940 |
| RUSC2 siRNA | Thermo Scientific | siRNA ID: s19070 |
| Podxl siRNA | Thermo Scientific | siRNA ID: s10771 |
| DENNd1a | Thermo Scientific | siRNA ID: s33637 |
| DENNd1b | Thermo Scientific | siRNA ID: s29140 |
| DENNd1c | Thermo Scientific | siRNA ID: s36719 |
| Cytodex Microcarrier Beads | Sigma-Aldrich | C3275-10G |
| CK-666 | Sigma-Aldrich | SML0006-5MG |
| Trimethoprim (TMP) | Sigma-Aldrich | T7883-5G |
| NP-G2-044 (Fascin Inhibitor) | Selleck Chem | S2962 |
| GGA3 PBD-Beads | Cytoskeleton, Inc. | GGA07-A |
| High Capacity Reverse Transcription Kit | ThermoFisher | 4368814 |
| Fibrinogen Type 1-S from Bovine Plasma | Sigma-Aldrich | F8630-1G |
| Thrombin from Bovine Plasma | Sigma-Aldrich | T7513-500UN |
| Aprotinin Protease Inhibitor | ThermoFisher | 78432 |
| Phenol-Red (Zebrafish Injection Mixture) | Avantor/ VWR | 34487-61-1 |
| CRIPSR gRNA | Integrated DNA Technologies (IDT) |  |
| Alt-R® S.p. Cas9 Nuclease V3, 100 µg | Integrated DNA Technologies (IDT) | 1081058 |
| CellTracker Deep Red | ThermoFisher | M22426 |
| 3-Aminobenzoic Acid Ethyl Ester (Tricaine) | Sigma-Aldrich | A5040-25G |
| Latex Beads, Polystyrene Carboxylate Mod | Sigma-Aldrich | L3280-1ML |
| Dynabeads™ Protein G for Immunoprecipitation | ThermoFisher | 10003D |
| MitoTracker DeepRed | ThermoFisher | M22426 |
| Trizol Reagent | ThermoFisher | 15596026 |
| Chloroform | Sigma-Aldrich | 288306 |
| MEGAscript™ T3 Transcription Kit | ThermoFisher | AM1338 |
| BCA Protein Assay Kit | ThermoFisher | 23225 |
| NHLF | Lonza | CC-2512 |
| HEK 293-A | ThermoFisher | R70507 |
| Microcarrier beads | Amersham | 17-0485-01 |
| Protease inhibitor cocktail | GoldBio | GB-334-20 |
| Agarose Resin | GoldBio | G-250-G |
| Fura Red™, AM, cell permeant | ThermoFisher | F3020 |

## ANTIBODIES

| Target Antigen | Vendor or Source | Catalog No./ Clone | Working Concentration |
| --- | --- | --- | --- |
| Rab35 | ThermoFisher | PA531674 | 1:500 (WB) |
| ACAP2 | ThermoFisher | PA557069 | 1:500 (WB) |
| OCRL | ThermoFisher | PA527844 | 1:200 (WB) |
| MICAL-L1 | ThermoFisher | PA5107177 | 1:200 (WB) |
| RUSC2 | ThermoFisher | PA572752 | 1:200 (WB) |
| Arf6 | Santa Cruz | sc-7971 | 1:200 (WB) |
| Myc-tag | ThermoFisher | 132500 | 1:1000 (IHC) |
| HA-tag | ThermoFisher | 26183 | 1:1000 (IHC) |
| cyan | Bio-Rad | AHP2986 | 1:1000 (IHC) |
| Alpha-tubulin | Abcam | ab52866 | 0.0648ug/mL<br>(1:10,000) (WB) |
| GAPDH | ThermoFisher | PA1988 | 1:1000 (WB) |
| Moesin | Abcam | ab52490 | 0.05ug/mL<br>(1:1000) (IHC) |
| VE-Cadherin | ThermoFisher | 14-1441-82 | 0.5ug/mL<br>(1:1000) (IHC) |
| Podocalyxin | R&D | AF1658 | 15ug/mL (1:200)<br>(WB & IHC) |
| Von Willebrand<br>Factor | Abcam | ab6994 | 10ug/mL<br>(1:1000) (IHC) |
| β1-Integrin | Abcam | ab30394 | 1:500 (IHC) |
| Phosphorylated<br>TIE-2/TEK<br>(Tyr992) | Sigma Aldrich | ABF131 | 0.25 ug/mL<br>(1:500) (IHC) |
| Anti-HA-Tag,<br>Rabbit<br>Monoclonal | Sigma-Aldrich | SAB5600116-100UG | 5ug/mL |
| Alexa Fluor™<br>488 Phalloidin | ThermoFisher | A12379 | 1 uM (1:200) |
| Alexa Fluor™<br>647 Phalloidin | ThermoFisher | A22287 | 1 uM (1:200) |
| Alexa Fluor™<br>555 Phalloidin | ThermoFisher | A34055 | 1 uM (1:200) |
| Goat anti-Rabbit<br>IgG (H+L)<br>Secondary<br>Antibody, Alexa<br>Fluor 488 | ThermoFisher | A11008 | 1ug/mL (1:500) |
| Donkey anti-Rabbit IgG (H+L)<br>Secondary<br>Antibody, Alexa<br>Fluor 555 | ThermoFisher | A31572 | 1ug/mL (1:500) |
| Donkey anti-goat<br>IgG (H+L)<br>Secondary<br>Antibody, Alexa<br>Flour 488 | ThermoFisher | A11055 | 1ug/mL (1:500) |
| Donkey anti-Goat IgG (H+L) Cross-Adsorbed Secondary Antibody, Alexa Fluor 555 | ThermoFisher | A21432 | 1ug/mL (1:500) |
| Chicken anti-Rabbit IgG (H+L) Cross-Adsorbed Secondary Antibody, Alexa Fluor 647 | ThermoFisher | A21443 | 1ug/mL (1:500) |
| Goat Anti-Rabbit HRP | Genesee Scientific | 20-303 | 1ug/mL (1:500) |

## OLIGOS AND SGRNA

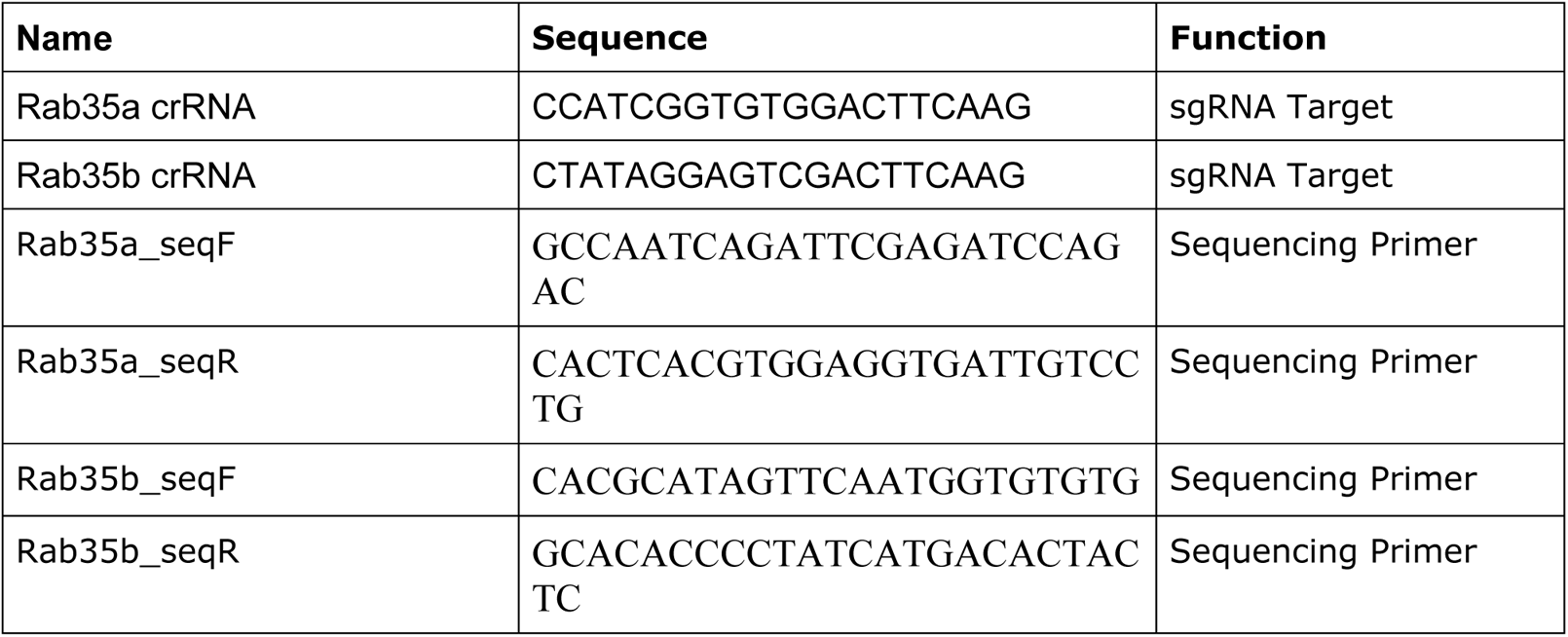

